# Sticker number modulates pyrenoid condensate assembly to support algal fitness

**DOI:** 10.64898/2026.01.27.701992

**Authors:** Gaurav Kumar, James Barrett, Philippe Van der Stappen, Alex Payne-Dwyer, Onyou Nam, Jack Shepherd, Lewis Frame, Jamieson Howard, Vuk Malis, Clément Dégut, Michael R. Hodgkinson, Ralf P. Richter, Charley Schaefer, Michael J. Plevin, Maria Hondele, Mark C. Leake, Benjamin D. Engel, Luke C.M. Mackinder

## Abstract

The valency of intrinsically disordered proteins underpins liquid-liquid phase separation (LLPS), yet how this parameter shapes condensate function and cellular fitness remains poorly understood. Here we exploit the algal pyrenoid–a minimal, two component LLPS system–to directly link condensate properties to physiological performance. Pyrenoid assembly is driven by a disordered, multivalent Linker protein that binds Rubisco at symmetry-related surface sites, with the number of binding motifs (“stickers”) varying across species. Using *Chlamydomonas reinhardtii*, we systematically tuned sticker number from two to nine and examined effects on Rubisco condensation, pyrenoid architecture and CO_2_ fixation. Three stickers were sufficient for condensation in vitro, but at least four were required for pyrenoid assembly in vivo. Cryo-electron tomography and single-molecule tracking revealed that increasing sticker number enhances Rubisco packing and mobility, while time-resolved imaging and competition assays demonstrated that sticker number governs the kinetics of pyrenoid formation and determines cellular fitness under fluctuating carbon conditions. Our findings establish sticker number as an evolutionary tuning parameter that balances condensate formation, dynamics, and function, providing a quantitative framework for linking the molecular grammar of phase separation to biological fitness.

## Introduction

A fundamental mechanism of cellular organisation is the biomolecular condensation of macromolecules (*e.g.* protein and nucleic acids) driven by multivalent interactions between constituent components^1^. In many cases, these interactions give rise to liquid-liquid phase separation (LLPS), in which molecules demix into a condensed phase with liquid-like material properties^2–6^. A useful framework for understanding LLPS in some contexts is the stickers-and-spacers model, in which discrete adhesive “stickers” interact either homotypically (with the same type of macromolecule) or heterotypically (with other types of macromolecule), through reversible non-covalent bonds to drive condensation above a critical concentration^5,7–10^. Stickers are separated by spacer regions which may be structured surfaces of globular proteins, or intrinsically disordered regions (IDRs), that interact with the solvent and favour multivalent interactions between the stickers in the system^11–13^. The multivalency, or sticker number, of the component proteins is crucial for LLPS^14^.

While the role of sticker number in setting critical concentrations and phase boundaries is increasingly well defined^9,15^, far less is understood about how tuning the sticker number of a single component in a heterotypic system influences condensate structure and function. In particular, it remains unclear whether variation in sticker number alters packing density and material properties, and how such changes impact condensate function, cellular fitness, and survival.

The Rubisco-Linker protein condensates at the core of biophysical CO_2_-concentrating mechanisms (CCMs) provide an ideal system to address this gap. In diverse algae and cyanobacteria, multivalent Linker proteins crosslink Rubisco to form dense condensate microcompartments—pyrenoids in algae and carboxysomes in cyanobacteria—that enhance CO_2_ fixation efficiency by concentrating Rubisco with its substrate (Fig. 1a, Fig. S1a). These condensates collectively mediate ∼180 gigatons of CO_2_ fixation annually, approximately half of the global net primary production^16–18^. Despite their conserved function, Linker proteins show little primary sequence similarity and are thought to have evolved convergently^19–21^.

**Fig. 1:**
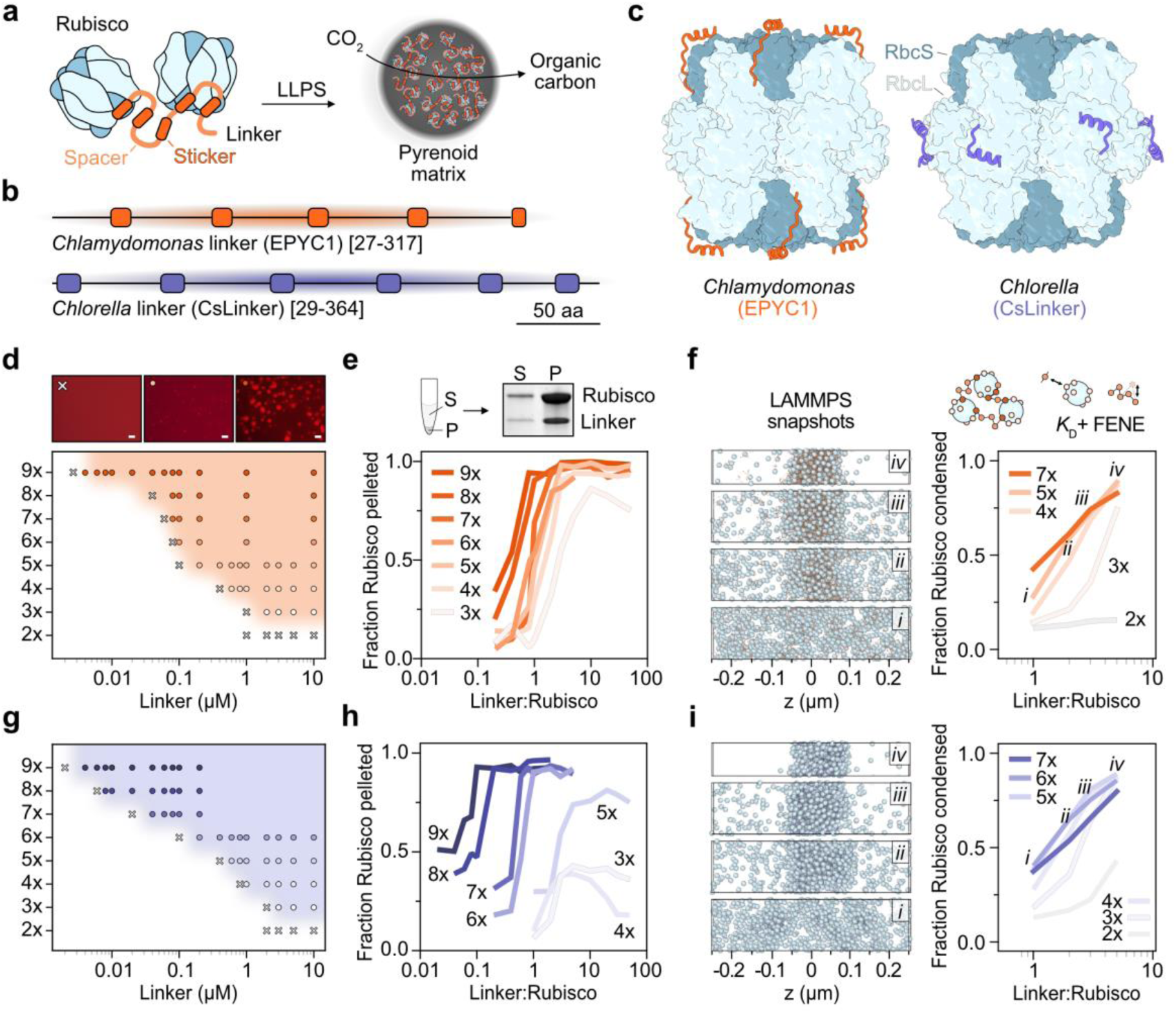
Linker protein sticker number dictates Rubisco phase separation critical points and partitioning. **a)** Schematic representation of Linker protein mediated Rubisco phase separation to form the pyrenoid matrix. **b)** Domain organisation of EPYC1 (orange) and CsLinker (purple), highlighting stickers (binding motifs; rectangles) and spacers (lines). **c)** Experimentally determined binding positions of EPYC1 and CsLinker stickers on Rubisco from PDB 7JN4 and 8Q05 respectively. **d)** Determination of sticker number-dependent *in vitro* phase separation critical points by microscopic observation of 1 µM CrRubisco (0.5% Atto594 labelled) with varying concentrations of Linker protein variants. Representative images show the critical points. Scale bar = 10 µm. **e)** Fraction of 1 µM CrRubisco recovered in the pelleted fraction following sedimentation assays with EPYC1 sticker variants. Points represent averages of two replicate experiments. **f)** Representative LAMMPS simulation snapshots (i-iv) from increasing concentrations of 5x EPYC1 against fixed CrRubisco, quantified in the indicated points adjacent. Plot shows mean partitioning across the final 100 frames of LAMMPS simulations across sticker number variants and concentrations. Points i-iv show quantifications of simulations with 5x EPYC1 at Linker:Rubisco ratios as indicated and shown in adjacent snapshots. **g)** Critical concentration of CsRubisco phase separation with CsLinker variants, as in d. **h)** Droplet sedimentation assays with CsRubisco and CsLinker variants, as in e. **i)** LAMMPS simulations with CsRubisco and CsLinker variants, as in f. Points i-iv show quantifications of simulations with 6x CsLinker at Linker:Rubisco ratios as indicated and shown in adjacent snapshots.

Algal pyrenoids display hallmark features of heterotypic LLPS^22^: they can dissolve and recondense in response to environmental cues, exhibit rapid internal mixing, and show liquid-like rearrangements on physiological timescales. These dynamic properties are particularly important given that algae experience fluctuating light, CO_2_, and temperature in their aquatic environments. As such, pyrenoids are an ideal model for exploring how multivalency in disordered proteins influences LLPS, and how condensate assembly impacts functional and physiological outcomes, including CO_2_ fixation and cell growth. Pyrenoids are also sufficiently large for detailed fluorescence microscopy and contain Rubisco structures resolvable by cryo-electron tomography (cryo-ET), enabling multi-scale analysis from *in vivo* dynamics to single macromolecular organization within the condensate^22,23^.

Conceptually pyrenoid assembly can be described within the “stickers-and-spacers” framework^11^: Linker proteins contain sequence-defined adhesive motifs (“stickers”), also termed Rubisco-binding motifs (RBMs), which mediate multivalent interactions with defined surface patches on Rubisco (Fig. 1b,c)^18,23,24^. For example, in *Chlamydomonas reinhardtii* (Cr), the Linker protein EPYC1 contains multiple stickers that each may interact with one of the eight small subunits of CrRubisco^23^. Linker-protein-mediated crosslinking is also observed in cyanobacterial carboxysomes, where Linker functionality is embedded within larger multi-domain proteins, highlighting the broader relevance of this multivalency-based mechanism (Fig. S1c)^25,26^.

In this study, we investigate how variation in sticker number influences Rubisco phase separation, and condensate assembly, and function, using the algal pyrenoid as a model system. Using a combination of *in vitro* biochemical assays and computational modelling, we examine how sticker number, Linker protein concentration, and sticker binding positions on Rubisco govern condensate formation. We then extend our investigation *in vivo* in *Chlamydomonas reinhardtii* to assess how sticker number modulates pyrenoid assembly, Rubisco partitioning, Rubisco packaging density, CO_2_-fixation efficiency, and cellular fitness. Collectively, these studies reveal sticker number as a tunable parameter that links molecular-level multivalency to condensate functionality and cellular fitness.

## Results

Linker proteins appear to have evolved convergently, and although their overall sticker-spacer composition and general physicochemical features (*e.g.*, hydrophilicity and high isoelectric points) are broadly conserved, there is variation in both the number and sequence of stickers as well as in spacer length and composition (Fig. S1d). In parallel, while Rubisco structure is highly conserved, recent work has shown considerable diversity in the positions of Linker binding sites on the Rubisco surface^23,25–27^ (Fig. S1e). To examine how these factors relate, we quantified sticker variation across the Rubisco-binding regions of the four Linker proteins known to undergo obligate heterotypic phase separation with their partner Rubiscos. Across >500 unique CsoS2 and CcmM35 homologs, we found that CsoS2 almost exclusively contains 4 stickers, whereas CcmM35 sequences typically contain 3 or 4. Similarly, analysis of EPYC1 and CsLinker homologs revealed a preference for 5 or 6 stickers, though with fewer representative sequences (Fig. S1f). Thus, while sticker number can vary, it is strongly clustered within each Linker family. This convergence suggests that sticker number, together with binding site geometry and interaction strength, may be a key evolutionary parameter controlling Rubisco phase separation and raising the central question of how sticker number influences the functionality of Rubisco in the condensate.

To study the importance of sticker number on Rubisco phase separation in pyrenoids, we selected the *Chlamydomonas reinhardtii* Linker protein, EPYC1, and the *Chlorella sorokiniana* Linker protein, CsLinker as well-characterised models. While both are highly hydrophilic, positively charged, and use short helical stickers to bind their respective Rubiscos (Fig. 1c, Fig. S1c,e), they differ in sticker number, sequence, binding affinity and binding geometry (Fig. 1b,c, Tables S1-3). EPYC1 has five stickers and interacts with the small subunits of *Chlamydomonas* Rubisco (CrRubisco) in a polar configuration with an apparent single sticker affinity of 414 µM^13^, while CsLinker has one sticker more (6 stickers) and binds equatorially to the large subunits of CsRubisco with an affinity of 100 µM (Fig. 1c)^27^.

### Sticker number modulates critical concentration and partitioning efficiency

To analyse the effect of sticker number on Rubisco phase separation *in vitro*, we recombinantly produced variants of EPYC1 and CsLinker comprising 2-9 repeating units (2x–9x). We centred the repeating units on the characterised Rubisco-binding domains such that each unit contained half of the preceding and following spacer, and units were truncated or expanded from the wild type (WT) sequence (see methods) (Fig. S2, Tables S1-3). We initially tested the effect of sticker number on the critical concentration of *in vitro* condensate formation by titrating the variants against a fixed Rubisco concentration of 1 µM, which we purified natively from each species. For both Linker proteins, at least three stickers were required to phase separate Rubisco, with 3x EPYC1 and 3x CsLinker variants forming visible droplets at concentrations of 2 and 3 µM, respectively, whereas 2x variants failed even at 50 µM, the maximum concentration tested (Fig. 1d, Fig. S3,S4). This indicates that two sticker variants of EPYC1 and CsLinker are insufficient to cross-link Rubisco molecules at physiologically relevant concentrations. For variants with >2 stickers, increasing the number of stickers per chain decreased both the critical chain concentration and the critical sticker concentration required to phase separate 1 µM Rubisco *in vitro* (Fig. 1d,g, Fig. S5). Strikingly, variants that matched the WT sticker number (5x EPYC1 and 6x CsLinker) showed nearly identical fitted critical concentrations between 0.1–0.2 µM (Fig. S5). Using 9x variants of both EPYC1 and CsLinker, we observed sub-micron droplet formation at concentrations as low as 6 and 4 nM respectively (Fig. 1d,g, Fig. S3,S4), consistent with the ability of longer, higher valency Linker proteins to cross-link more Rubisco molecules per chain. The observed droplet size was also correlated with linker and sticker concentration, suggesting that Rubisco partitioning in the condensed phase is proportional to Linker protein concentration. These observations motivated us to examine more precisely how Linker protein concentration influences Rubisco partitioning in the condensed phase.

To quantify the amount of Rubisco partitioned into droplets, we performed titration sedimentation assays with 3x-9x variants of EPYC1 and CsLinker, and made two observations that hold true for both EPYC1 and CsLinker (Fig. 1e,h). First, Rubisco partitioning into the condensed phase increased with Linker protein concentration and plateaued at higher concentrations, indicating that partitioning is concentration-dependent regardless of sticker number and consistent with the droplet size changes observed in Fig. 1d,g. Second, at a given Linker protein concentration, Rubisco sedimentation increased with sticker number, suggesting that higher sticker numbers promote greater cross-linking of Rubisco. Notably, at sticker numbers >5, CsLinker variants required substantially lower concentrations to sediment >90% of Rubisco compared to equivalent EPYC1 variants, whereas at sticker numbers <5, Rubisco sedimentation was greater for EPYC1 than for the corresponding CsLinker variants (Fig. 1e,h). Interestingly, the sticker number variants corresponding to the WT sequences (*i.e.* 5x for EPYC1 and 6x for CsLinker) shared a similar partitioning saturation concentration at ∼2x the Rubisco concentration, consistent with *in vivo* observations^27,28^.

In parallel, we tested the predictive power of an updated statistical physics approach^13^, and developments to state-of-the-art Large-scale Atomic/Molecular Massively Parallel Simulator (LAMMPS) simulations^29^ for modelling Rubisco:Linker condensation in the sticker-and-spacer framework^11^. In our statistical physics approach, where we use Rubisco dimerization as a proxy for critical concentration, we parameterize the interaction between spherical Rubisco particles, with binding sites at their experimentally determined positions, and the Linker protein using a small number of parameters (K_D_, the sticker binding strength, and the spacer flexibility (*l*_K_). Although previous predictions where *Chlamydomonas* Rubisco was modelled as a cube showed good predictive power for 3-5 stickers^13^, we found an excluded volume effect of the spherical model to drastically impact our prediction of the *Chlamydomonas* system, resulting in increased binding energy (11.8 vs 12.5 *k*_B_*T* respectively) and decreased spacer flexibility (0.88 vs 0.58 nm) (Fig. S5). New measurements of EPYC1 using atomic force microscopy (AFM) indicated a spacer flexibility of EPYC1 within the range of the two models (*l*_K_ 0.72 ± 0.09 nm S.D.) (Fig. S6), indicating further parameterization of the model in this geometry is required. Although we could not uniquely parameterize our model to *Chlorella* biophysical data, we observed another sizable shift to higher dimerization concentrations when the binding sites were positioned equatorially on Rubisco, as determined for CsLinker^27^ (Fig. S5), a similar effect as observed in Grandpre et al. (2023).

In our LAMMPS simulations, which are parameterized using similarly few parameters (the *K*_D_, the sticker binding strength through the well depth, and the spacer flexibility through the FENE spring constant), where Rubisco is also modelled as a sphere, we monitored Rubisco condensation as a function of sticker number and linker concentration. Although the simulations were broadly consistent with experiments, showing increased Rubisco partitioning with both sticker number and concentration in both systems (Fig. 1f,i), they failed to capture two nuances of our *in vitro* data. First, in experiments, CsLinker variants with 3x and 4x stickers failed to partition more than 50% of Rubisco into the condensed phase, whereas this limitation was not observed in simulations. Second, the pronounced effect of increasing sticker number on CsLinker partitioning seen experimentally was absent in simulations. It appears the strong influence of adding a sixth sticker to CsLinker cannot be captured with the current simple stickers-and-spacer framework that includes simple descriptions for sticker binding, spacer extensibility, and Rubisco shape. Thus, models to predict subtle changes in the system require more detailed molecular (or even atomistic) resolution of linker and Rubisco properties.

### A sticker number threshold determines Rubisco partitioning and density in the pyrenoid

Building on our understanding of how sticker number affects Rubisco condensation, we next examined these effects *in vivo* and their impact on pyrenoid formation, function, and cellular fitness. We used the tractable *Chlamydomonas* pyrenoid system, where deletion of the native Linker protein EPYC1 disrupts pyrenoid assembly and impairs growth^18^. To visualize Rubisco condensation, we generated a “chassis” line in an *epyc1* knockout background that expressed an extra copy of RbcS fused to mVenus, such that 0.95% of the RbcS in the total Rubisco pool was fused to mVenus (*i.e.* for every 13 Rubisco holoenzymes, only one mVenus tag is present) (Fig. S9). This chassis line expressed Rubisco at WT levels but did not condense Rubisco to form the pyrenoid matrix (Fig. 2b, Fig. S10c). We used a recently developed chloroplast expression toolkit^30^ to express EPYC1 sticker variants (2x-9x) from a neutral insertion locus in the chassis, resulting in equal expression of the variants across lines, which was comparable to the expression of EPYC1 in the WT background (Fig. S10).

**Fig. 2:**
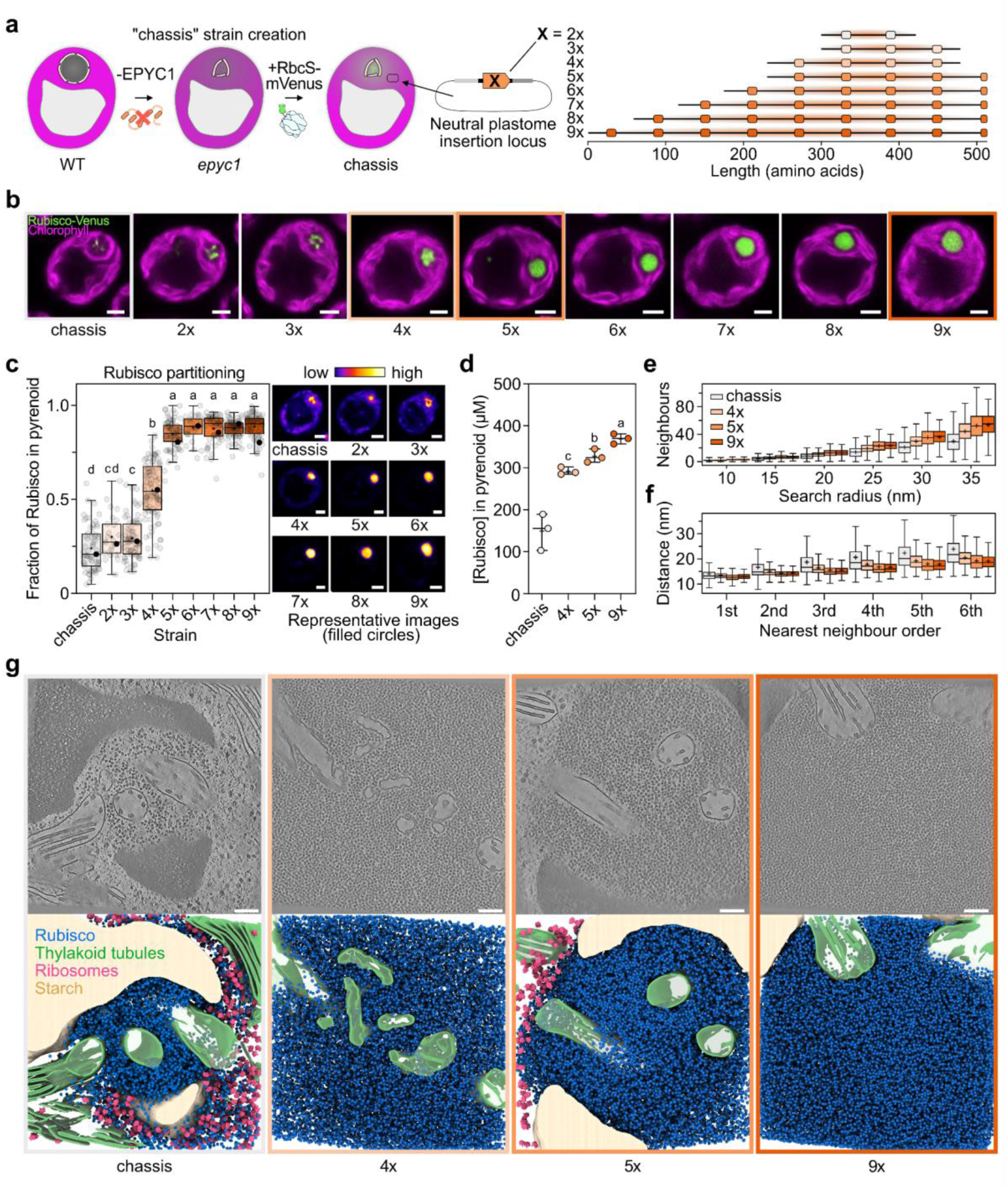
Linker protein sticker number tunes Rubisco partitioning and crowding in pyrenoids. **a)** Generation of the chassis strain from the *epyc1* knockout background^18^, by introduction of an additional copy of RbcS fused to the mVenus fluorophore. EPYC1 sticker variants were subsequently introduced from a neutral locus in the chloroplast genome (plastome). **b)** Representative confocal microscopy images illustrating the morphology of Rubisco-mVenus signal in the pyrenoid of cell lines after 18 hours of photoautotrophic growth. Scale bar = 1 µm. **c)** Quantification of Rubisco-mVenus partitioning at the canonical pyrenoid position of each line. Box plots indicate the median (line), mean (cross), IQRs (box) and outlier range (whiskers, 1.5x IQR). Filled circles represent quantifications in representative images adjacent. Scale bar = 1 µm. **d)** Rubisco concentration within pyrenoids based on cryo-ET analysis. Median (line), mean (cross) and range shown. One-way ANOVA with post-hoc Tukey test, p < 0.05 grouping. **e)** Neighbour number analysis from template matching. **f)** Nearest neighbour distance analysis. Box plots as in c. **g)** Representative slices through cryo-ET volumes (top) and 3D reconstructions (bottom) of pyrenoids from chassis, 4x, 5x, and 9x strains. Rubisco (blue), tubules (green), starch (light brown) and ribosomes (pink) are shown. Scale bars = 100 nm.

We first examined these lines by fluorescence microscopy 18 hours after switching the cultures from mixotrophic (supplemented with acetate), to phototrophic (without acetate) growth conditions under ambient CO_2_ (∼420 ppm); conditions under which the CCM is fully induced^31^, and the pyrenoid matrix is fully condensed at the canonical position^32^. Lines expressing 2x or 3x EPYC1 variants did not form a pyrenoid matrix, though did display some signal at the canonical pyrenoid position, suggestive of a rudimentary Rubisco network. The 4x variant produced visible Rubisco condensation, though with a significantly less circular cross-section, with median roundness of 0.83 (IQR: 0.75-0.89, *p* < 0.0001) than seen in lines expressing 5x-9x variants, which all had median roundness values between 0.89-0.94 (IQR: 0.84-0.96) (Fig. 2b, Fig. S11c). Rubisco partitioning into the condensate was quantified from mid-cell sections by measuring RbcS-mVenus signal within the segmented pyrenoid region, following Barrett et al. (2024). Across strains, the extent of partitioning scaled with pyrenoid size (Fig. S11d,e). Cells expressing 2x and 3x variants showed median partitioning values of 27%, similar to the 21% partitioning observed in the chassis. By contrast, the 4x variant exhibited a wide spread of partitioning values with a median of 55% (IQR: 44-68%), consistent with its irregular morphology. Variants with 5-9 stickers had median partitioning values of >80% with a much tighter spread of values (IQR of 5x-9x: 80-93%) (Fig. 2c), indicating that condensation becomes highly efficient at five stickers, with additional stickers providing little further increase. These results suggest that ∼5 stickers represent a threshold for robust and effective Rubisco condensation and recruitment into the pyrenoid.

Motivated by the clear sticker number-dependent changes in Rubisco partitioning, we next asked whether the variation in sticker number might lead to differences in the structural organization of Rubisco in the pyrenoid. We utilised cryo-electron tomography (cryo-ET), which has recently become a powerful tool to study biomolecular condensates both *in vitro* and *in v*ivo^33–35^. We collected three high-quality tomograms each for the chassis, 4x, 5x, and 9x lines, all grown to mid-exponential phase under phototrophic conditions at ambient CO_2_ (∼120 h). These lines were selected to represent key benchmarks: the chassis as a no-condensation control; the 4x line as the threshold with irregular and variable condensates; the 5x line as the WT sticker number, and the 9x line as the maximum sticker number tested. Consistent with our fluorescence microscopy observations, cryo-ET revealed clear pyrenoid matrix formation in the 4x, 5x and 9x lines, while the chassis line contained a smaller, less dense Rubisco condensate (Fig. 2g, Movie S1). Despite lacking a fully developed pyrenoid matrix, the Rubisco condensate in the chassis line was positioned at the canonical location, intersected by reticulated thylakoids and surrounded by a starch sheath, suggesting the rudimentary Rubisco network observed in the absence of EPYC1 may be mediated by membrane- or starch-associated Rubisco binding proteins present at the canonical pyrenoid position^36–38^. Interestingly, when the chassis line was grown mixotrophically in acetate-supplemented media, although still highly reduced when compared to the 4x, 5x and 9x lines, the Rubisco condensate was larger and more densely packed than in obligate phototrophic conditions (Fig. S12), suggesting unexplored changes in Rubisco or Rubisco-tethering protein expression may contribute.

Using high-resolution template matching inside precise matrix masks (Fig. S13), we found that Rubisco packing density and effective concentration in the pyrenoid matrix increased with sticker number. The chassis exhibited the lowest concentration (149 ± 25 µM s.e.m.), while Rubisco concentration in the 9x variant (369 ± 7 µM) was 26% more densely packed than the 4x line (291 ± 5 µM) and 12% denser than the 5x line (328 ± 9 µM) (Fig. 2d). Interestingly, this trend was also recapitulated in our LAMMPS simulations (Fig. S7e). Nearest-neighbour analysis of the cryo-ET data confirmed the positive correlation between sticker number and Rubisco concentration, with Rubisco complexes most sparsely distributed in the chassis and progressively closer together from 4x to 9x (Fig. 2e,f). However, the first neighbour positioning was nearly identical in all lines (median center-to-center distance = 13.2, 13.2, 12.7, 13.0 nm for the chassis, 4x, 5x, and 9x, respectively), also matching measurements of WT *Chlamydomonas* pyrenoids from other recent cryo-ET studies^39,40^. This indicates that there is a preferred minimum distance between Rubisco complexes, which may be set by the Linker proteins that cross-link them. Rather than reduce the minimum distance between neighbouring Rubisco, pyrenoids containing longer EPYC1 variants with higher sticker number have fewer unoccupied spaces in their matrix, perhaps due to additional Rubisco cross-linking.

Comparing the fluorescence microscopy and cryo-ET measurements, it is apparent that while Rubisco partitioning into the condensate and its packing density within the matrix both increase with sticker number, these two properties do not scale identically. For example, in the 4x line, the fraction of Rubisco localized to the pyrenoid was substantially lower than in the 5x line (Fig. 2c), yet the internal packing density differed only modestly (Fig. 2d). Conversely, although the 5x and 9x variants showed similar partitioning at the time of imaging, the 9x pyrenoid displayed noticeably higher packing density. However, taken together, these observations suggest that sticker number in Linker proteins is critical for optimizing Rubisco partitioning and crowding in the pyrenoid condensate.

### Sticker number modulates Rubisco mobility

Given the clear differences we observed in partitioning and packing density between sticker variants, we hypothesised that Rubisco dynamics within the matrix may also be affected. We used fluorescence recovery after photobleaching (FRAP) to assess internal mixing in the pyrenoid, using the Rubisco-mVenus signal as described previously^22^ (Fig. 3a, Fig. S14). We evaluated all sticker variants using lines grown under phototrophic conditions for 12 h under ambient CO_2_. Re-homogenization of the Rubisco-mVenus signal following photobleaching occurred with a half-life of ∼20 s in all lines (Fig. 3b,c). Surprisingly measured half-life values did not differ significantly between the sticker variant lines tested, suggesting that ensemble Rubisco rearrangement is not influenced by sticker number. To explore dynamics at higher temporal and spatial resolution, we tracked Rubisco-mVenus mobility with single molecule precision using SlimVar microscopy^41^ in the 4x, 5x and 9x variant cell lines grown under the same conditions we used for the cryo-ET experiments (∼120 h in phototrophic conditions) (Fig. 3d). We observed that Rubisco in the 5x and 9x lines had similar median diffusivities of 0.4 µm^2^ s^-1^ (IQR: 0.008-1.527 and 0.03-1.384 µm^2^ s^-1^ respectively). In contrast, the 4x line showed significantly slower diffusion (0.16 µm^2^ s^-1^; IQR 0.0002-0.98; *p* < 0.0001) (Fig. 3e). We speculate that this difference arises in part from a larger fraction of relatively immobile Rubisco molecules falling below the experimental detection limit (0.08 µm^2^ s^-1^), which was 43% in the 4x line compared to 30% in the 5x and 9x lines (Fig. 3e). To validate this observation, we conducted SlimVar assays using condensates reconstituted from recombinantly expressed 4x, 5x, and 9x Linker proteins and the Rubisco / Rubisco–mVenus pool that we purified from the chassis line. Although diffusivities were substantially higher than those measured *in vivo*, the same relative trend was observed. Median Rubisco-mVenus diffusivity in 5x and 9x condensates was similar (1.1 µm^2^ s^-1^; IQR 0.34-2.63 and 0.35-2.67 µm^2^ s^-1^ respectively) and significantly faster than in 4x condensates (0.83 µm^2^ s^-1^; IQR: 0.26-2.17; *p* < 0.0001) (Fig. 3f). Only a small fraction of molecules were below the experimental sensitivity (0.03 µm^2^ s^-1^) with comparable proportions across the samples (4.9% in 4x, 4.2% in 5x and 3.5% 9x). Our tracking analysis also detected transient particles containing one or more Rubisco-mVenus molecules moving within the dense phase. This observation shows that effective unbinding of Linker proteins from cross-linked Rubiscos is slower than diffusion within the condensate, as each region of ∼40 nm diameter took longer to fully exchange Rubisco-mVenus signal than the average track lifetime (35 ± 5 ms). Together, the cryo-ET and single-molecule tracking data indicate that the 4x line exhibits reduced Rubisco mobility and forms a less densely packed condensate.

**Fig. 3:**
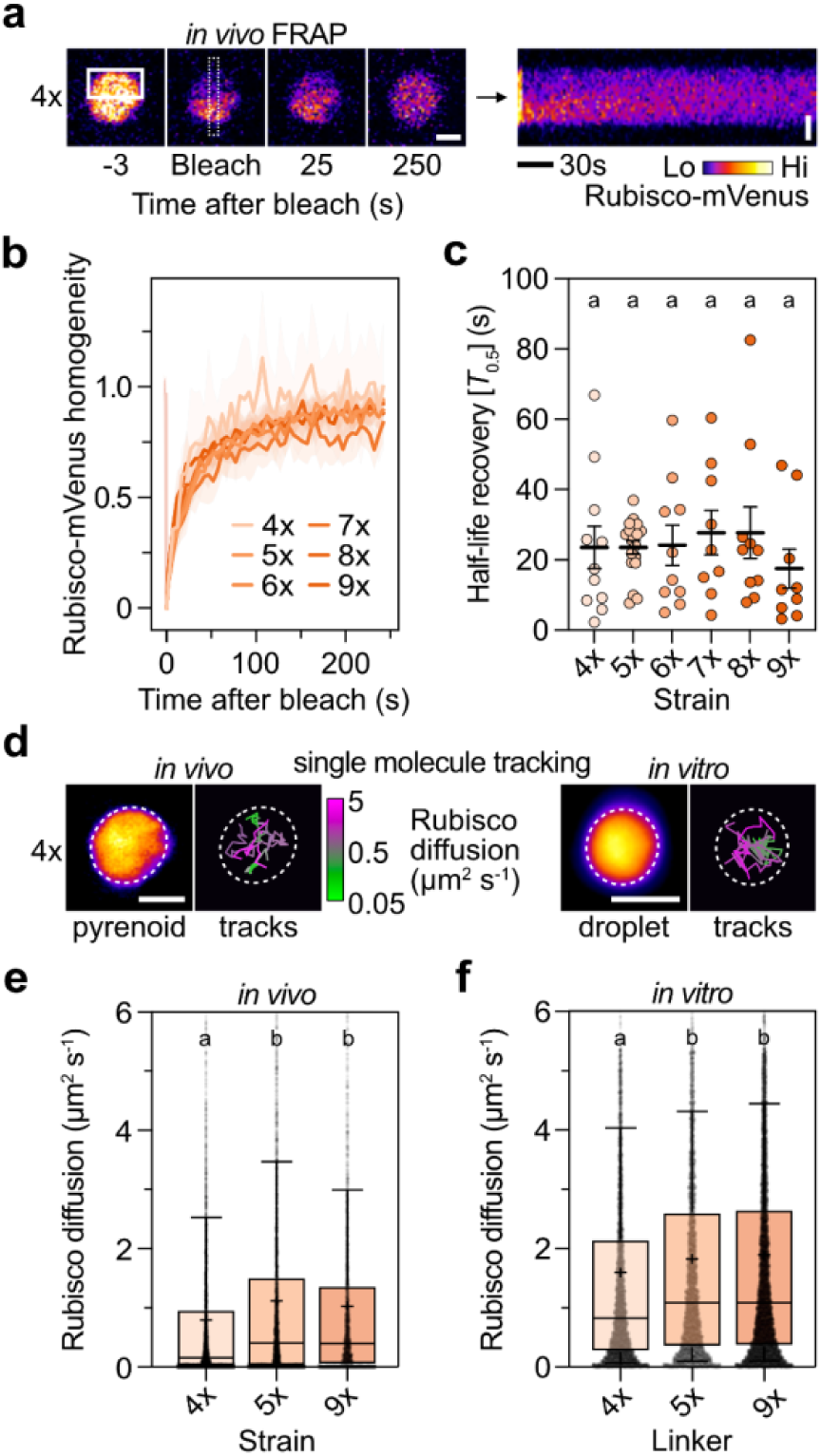
Sticker number modulates Rubisco mobility. **a)** Example fluorescence recovery after photobleaching (FRAP) experiment of Rubisco-mVenus in the pyrenoid of the 4x line. The solid white line indicates the bleach region, the dashed line indicates the region used for homogeneity analysis as in the adjacent kymograph. Scale bar = 1 µm. **b)** FRAP homogeneity recovery curves. Lines represent the mean of replicates, with the shaded region representing the s.e.m. **c)** Fitted *T*_0.5_ values from individual replicates, where the mean and s.e.m. are represented by the lines and error bars. **d)** Examples of Rubisco-mVenus single molecule tracking *in vivo* in the pyrenoid of the 4x line and *in vitro* in droplets formed with the 4x protein. The dashed line denotes the pyrenoid/droplet boundary, with example tracks shown within, coloured according to their rate of diffusion. Scale bar = 1 µm. **e)** Rubisco diffusivity in the pyrenoid matrix *in vivo*. Box plots indicate the median (line), mean (cross), IQRs (box) and outlier range (whiskers, 1.5x IQR). Statistical groups have *p* < 0.0001. **f)** Rubisco diffusivity in *in vitro* droplets formed with Rubisco / Rubisco-mVenus purified from the chassis. Box plots and statistical significance as in e.

### Sticker number governs the temporal dynamics of Rubisco condensation

The apparent discrepancies in Rubisco partitioning, density, and mobility between the 4x, 5x and 9x lines prompted us to systematically investigate the changes in Rubisco partitioning over time following induction of pyrenoid matrix formation, following the switch from mixotrophic to phototrophic growth conditions. Cells were grown to mid-exponential phase in acetate-supplemented media (mixotrophic), where Rubisco is partially diffuse in the chloroplast stroma^32^, before inducing pyrenoid formation and CCM activation by switching to acetate-free media under ambient CO_2_ conditions. At the 0 h timepoint, 5x and 9x lines already partitioned most of their Rubisco in the pyrenoid (76%; IQR 66-83% and 78%; IQR 66-90%) whereas 4x exhibited much lower partitioning (50%; IQR 43-66%) (Fig. 4a,b), reflecting the pattern we observed after 18 h (Fig. 2c). We monitored partitioning over the subsequent 100 h of exponential growth, during which all lines reached similar near-maximal partitioning (4x: 91%; IQR 88-94%, 5x: 91%; IQR 89-93%, 9x: 91%; IQR 88-93%). Despite similar endpoints, the dynamics of partitioning differed markedly. The 9x line reached maximal partitioning fastest (12 h), while 5x partitioned more rapidly than 4x but had not plateaued by 24 h (Fig. 4a,b). The most pronounced differences occurred during the 0-12 h period, when partitioning in the 4x line was significantly lower than in the 5x and 9x lines (*p* < 0.0001). We observed similar trends in time-resolved droplet sedimentation assays with the 4x, 5x, and 9x variants mixed with Rubisco. Consistent with in vivo measurements, Rubisco partitioning was slower in the 4x variant, while the 9x variant exhibited the fastest sedimentation, condensing all Rubisco within the first minute of mixing (Fig. 4c). While there is a clear disconnect between the timescales of partitioning *in vivo* vs. *in vitro*, we hypothesise that this physical effect is likely paired with some regulatory process (e.g., phosphorylation of EPYC1^42^) that together contribute to the sticker-dependent temporal dynamics of Rubisco condensation.

**Fig. 4:**
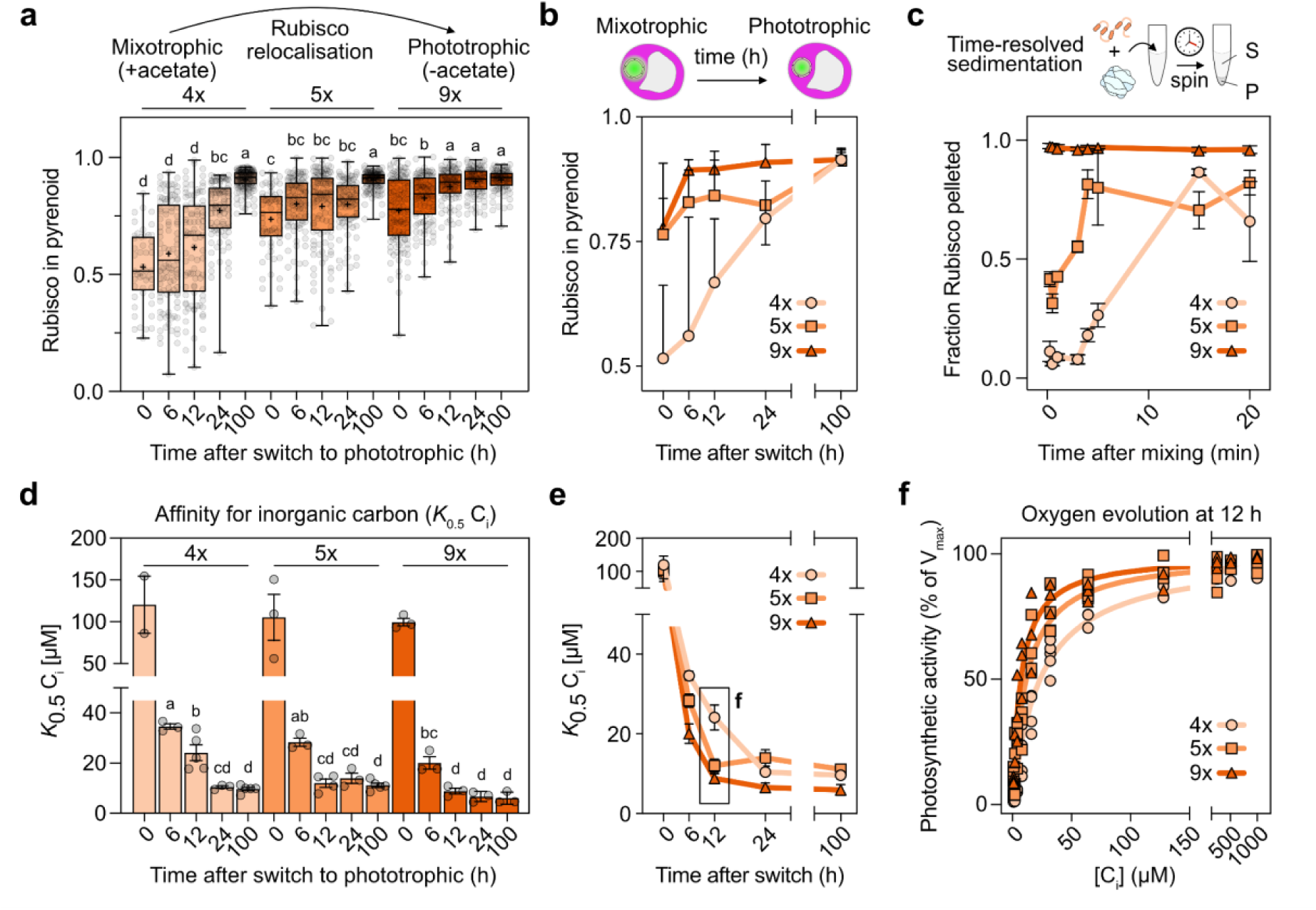
Sticker number is critical for timely Rubisco condensation and efficient CO_2_ fixation. **a)** Time-dependent Rubisco partitioning into the pyrenoid following the switch from mixotrophic to phototrophic growth conditions. Box plots indicate the median (line), mean (cross), IQRs (box) and outlier range (whiskers, 1.5x IQR). Statistical groups have *p* < 0.0001. **b)** Plot of median Rubisco partitioning against time, where error bars indicate IQRs. **c**) Time-dependent partitioning of Rubisco into *in vitro* droplets by time-resolved droplet sedimentation quantification. Mean ± s.e.m. are indicated. **d)** Affinity for inorganic carbon (*K*_0.5_ C_i_) measured by oxygen evolution. Mean ± s.e.m. are indicated, where statistical groups have *p* < 0.001. **e)** Plot of mean *K*_0.5_ C_i_ against time, where error bars indicate s.e.m. **f)** Oxygen evolution rate as a percentage of the fitted V_max_ as a function of inorganic carbon (C_i_) concentration, measured 12 hours after the switch from mixotrophic to phototrophic growth conditions. For 4x, 5x and 9x, n = 5, 4, and 3 respectively. The Michaeles-Menten fit to the average of the replicates is shown.

### Timely Rubisco condensation is essential for efficient CO_2_ fixation

Given the essential role of Rubisco condensation in cellular fitness^18^, we hypothesized that the differences observed in Rubisco partitioning dynamics following the switch from mixotrophic to phototrophic conditions would affect the efficiency of carbon fixation. To test this, we measured the photosynthetic affinity for inorganic carbon (*K*_0.5_ C_i_), using O_2_ evolution over the same time course as our partitioning experiments, after switching to phototrophic growth at ambient CO_2_ conditions. At 0 h, all lines exhibited high *K*_0.5_ C_i_ values, indicative of inefficient CCM function, consistent with both low Rubisco partitioning and the absence of induction of other essential CCM components^31^. By 6 h, the CCM is fully induced, so differences at later time points can be attributed more directly to Rubisco partitioning. Over the subsequent 100 h, the *K*_0.5_ C_i_ values of all lines decreased to a comparable minima (4x: 9.6 ± 0.7 µM s.e.m.; 5x: 11.1 ± 0.9 µM; 9x: 4.2 ± 0.4 µM), matching the value measured for WT at 100 h (11.0 ± 0.4 µM) (Fig. 4d,e, Fig. S16). Independent measurements after 100 h at 100 ppm CO_2_ (4x: 9.6 ± 0.7 µM; 5x: 11.1 ± 0.9 µM; WT: 11.0 ± 0.4 µM) (Fig. S16c,d), supported these observations and confirmed the chassis line remained inefficient (*K*_0.5_ C_i_: 131 ± 17 µM), consistent with its reduced Rubisco partitioning (Fig. 2b). The most striking observation emerged at 12 h, where the 4x line showed a significantly higher *K*_0.5_ C_i_ value (24.1 ± 3.2 µM, *p* < 0.001) than both the 5x and 9x lines, which had both reached near minimum values by this point (12.0 ± 1.7 µM and 8.8± 1.2 µM, respectively). This observation establishes a functional connection between sticker number of the Linker protein and early carbon-fixing efficiency of the pyrenoid, with reduced performance attributable to delayed or incomplete Rubisco partitioning during the onset of pyrenoid matrix condensation. These results indicate that while four EPYC1 stickers are sufficient for Rubisco condensation, a fifth sticker ensures timely condensation following CCM induction, enhancing the concentration and mobility of Rubisco molecules within the pyrenoid, necessary for efficient carbon fixation.

### Sticker number impacts cellular fitness

After extensively characterizing how sticker number influences Rubisco partitioning and the effect on affinity for inorganic carbon, we next examined whether these differences were reflected in the cellular fitness under different CO_2_ conditions. Under CO_2_-replete conditions (30,000 ppm), all sticker variant lines (2x-9x) and the chassis grew well in spot tests (Fig. 5a). By contrast, under ambient (∼420 ppm) and limiting (100 ppm) CO_2_ conditions, growth of the chassis and the 2x, and 3x lines was inhibited (Fig. 5a, Fig. S17), reflecting the lack of efficient Rubisco condensation into the pyrenoid matrix (Fig. 2b). Interestingly, despite slower Rubisco recruitment in the 4x variant, its growth under all conditions was comparable to the 5x-9x sticker variants and WT (Fig. 5a). These growth patterns were corroborated by liquid culture assays, which showed stunted growth in the chassis, 2x, and 3x lines, whereas lines with 4-9 stickers grew similarly to WT under both ambient and limiting CO_2_ conditions (Fig. 5b,c, Fig. S18). Together, these results suggest that partial Rubisco recruitment during pyrenoid biogenesis in the 4x line did not substantially compromise cellular fitness.

**Fig. 5:**
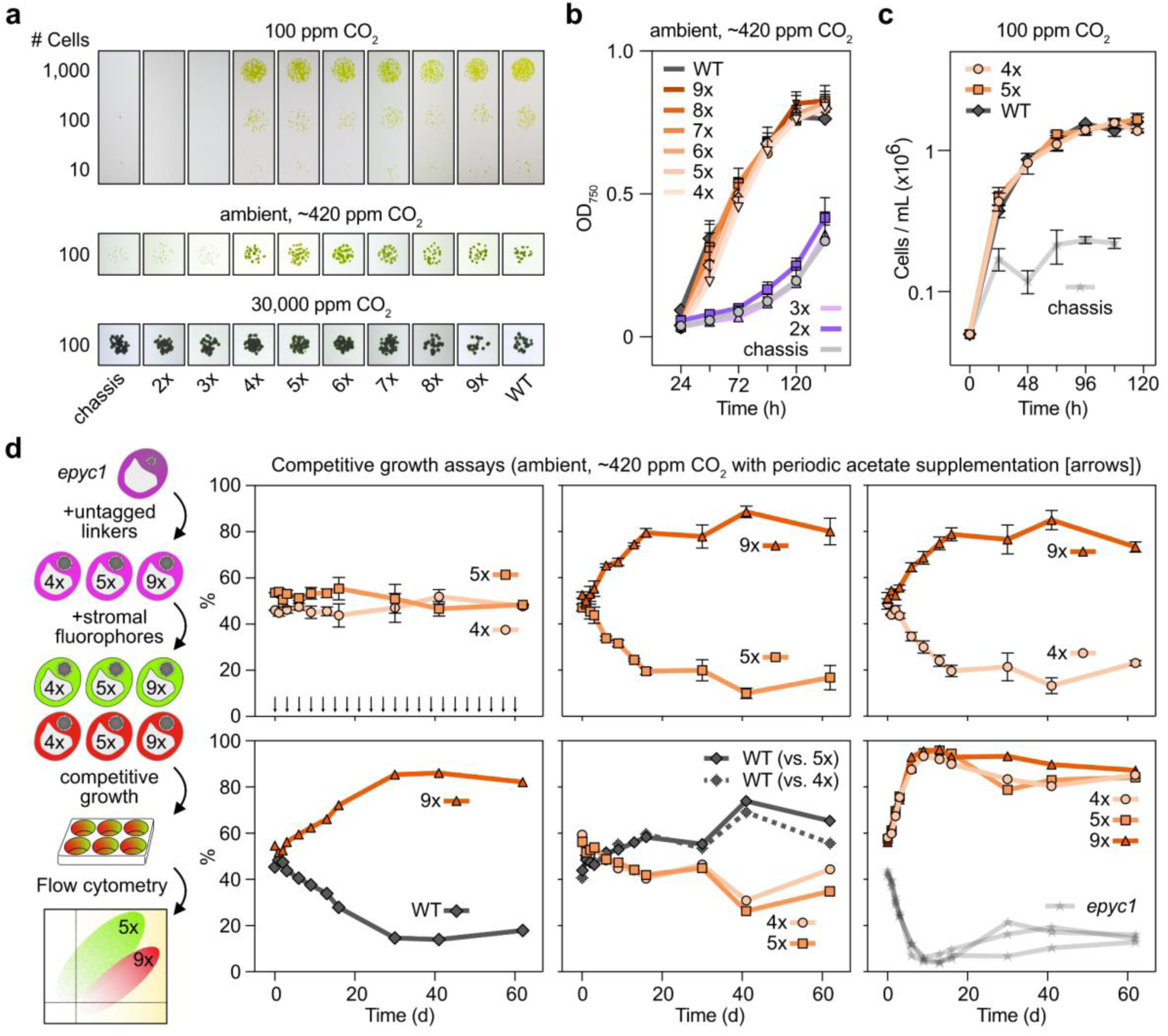
Sticker number impacts cellular fitness. **a)** Spot test growth assays of under limiting (100 ppm), ambient (420 ppm) and replete (30,000 ppm) CO_2_ conditions. **b)** Liquid growth assays of cells grown under ambient CO_2_ conditions. Mean ± s.e.m. is shown (n = 3). **c)** Liquid growth assays of cells grown under limiting CO_2_ conditions (100 ppm). Mean ± s.e.m. is shown (n = 3). **d)** Schematic of strain generation and workflow of competition growth assays (left) and proportion of each strain over time (right). The mean ± s.e.m. of the reciprocal line pairs is reported (see methods). Arrows indicate the points of dilution in 10% acetate-supplemented media.

Given the differences in C_i_ affinity we observed between the lines, we hypothesized that our growth assays were not sensitive enough to reveal subtle phenotypic differences between the strains. We therefore developed a flow-cytometry based competition growth assay to unequivocally distinguish growth differences between strains. To create lines for these experiments, we returned to the *epyc1* knockout background lacking Rubisco-mVenus in which we expressed the same untagged 4x, 5x and 9x EPYC1 variants as used throughout. Each subsequent strain was independently transformed with mVenus and mCherry markers, which were inserted at a separate neutral chloroplast locus and expressed to a low level in the chloroplast stroma. We then performed assays at conditions we expected to exacerbate phenotypic differences. At the onset, pairs of lines were mixed in equal proportions (*e.g.* 4x, mVenus with 5x, mCherry), and their reciprocal label swaps were also performed (*e.g.* 4x, mCherry with 5x, mVenus) to control for effects of the fluorescent markers. Because we had observed changes in Rubisco partitioning when shifting from mixotrophic (acetate - supplemented) to phototrophic (acetate-free) growth conditions, we supplemented cultures with acetate to a concentration 10% of that in acetate-supplemented media at periodic intervals. It has previously been shown that this concentration of acetate can be fully consumed within 24 h of growth^43^, and so we rationalised that periodic addition would trigger reduced Rubisco partitioning before being rapidly consumed and priming the competitive advantages between strains that we observed in the kinetics of Rubisco relocalisation to the pyrenoid (Fig. 4a,d). To this end we maintained the assays in exponential growth by dilution in 10% acetate media every 72 hours, and monitored population dynamics by flow cytometry, where we averaged the reciprocal experiments (Fig. 5d, Fig. S19, S20).

Surprisingly, we observed no substantial changes in the relative growth of the 4x and 5x lines in competition assays, whereas the 9x strain consistently outcompeted both 4x and 5x, as well as unlabelled WT cells (Fig. 5d). The 4x and 5x lines showed comparable fitness to WT, with neither displaying a conclusive competitive advantage. All sticker variant lines outcompeted the untagged *epyc1* knockout within the first seven days (Fig. 5d). These observations suggest that the 9x variant achieves enhanced CO_2_ fixation efficiency under changing metabolic conditions, likely due to faster or more effective Rubisco recruitment into the pyrenoid, as further supported by time-resolved confocal imaging. Moreover, increasing the number of stickers beyond 5, as in the 9x variant, provides a further advantage, likely by accelerating Rubisco recruitment or increasing Rubisco density, which enhances CO_2_ fixation efficiency under fluctuating conditions of CO_2_ availability. Collectively, these data demonstrate that the number of stickers in native Linker proteins directly impacts cellular fitness and that sticker numbers observed in nature have likely evolved to enable optimal condensate properties and hence cellular fitness across a range of selective conditions.

## Discussion

Phase separation has emerged as a general principle of cellular organization, enabling cells to dynamically compartmentalize biochemical reactions^44,45^, macromolecules^46^, and signalling processes^7^, without the need for membrane-bound organelles. A central challenge in this field is to understand how the condensate material properties are encoded by the sequence features of their components (*i.e.* stickers and spacers), and how those physical properties relate to biological function. Here, we used the algal pyrenoid as a model to link the molecular grammar of condensation, and the modulation of condensate material properties, directly to cellular physiology. The pyrenoid is uniquely positioned as a framework for testing how sticker number, affinity, and geometry together shape condensate formation, dynamics, and fitness, given its assembly around a defined multimeric enzyme, Rubisco, and an intrinsically disordered Linker protein, EPYC1.

We applied an integrated multi-scale approach to the pyrenoid system, combining in vitro reconstitution, statistical physics and coarse-grained modelling, single-molecule tracking, live-cell imaging, in situ cryo-ET, and physiological assays of carbon fixation and competitive growth. This enabled us to directly link EPYC1 valency to condensate architecture and cellular fitness, demonstrating how changes in Rubisco packing and mobility governed by sticker number underlie altered CCM efficiency and fitness under fluctuating CO_2_. These multi-scale measurements establish the pyrenoid as tractable model of phase separation in which quantitative predictions about condensate function can be evaluated against structural, cellular, and physiological readouts.

### Phase behavior emerges from a balance of valency, geometry, and affinity

By integrating *in vitro* experiments (Fig. 1), statistical physics approaches (Fig. S8), and coarse-grained molecular dynamics simulations (Fig. 1f,i), we explored how sticker number, binding affinity, and binding geometry collectively determine emergent phase behavior. Consistent with earlier theoretical work^29^, our extension of a recent statistical physics framework^13^ demonstrates that the spatial arrangement of sticker binding sites on the Rubisco surface exerts a strong influence on cross-linking efficiency (Fig. S8). This is especially notable given the striking lack of conservation in binding geometries across Rubisco–Linker systems^23,25–27,47^. Although it is impossible to completely decouple this effect from binding affinity changes in experiment, the combination of our *in vitro* and molecular dynamics results strongly suggest that binding site affinity and valency are balancing differences in binding site geometry, and in sum determining phase behaviours. The apparently lower cross-linking efficiency of the equatorial arrangement of CsLinker binding sites could provide a plausible explanation for the increased affinity of the individual sites and for the additional sticker in the chain. These factors, along with unexplored contributions from spacer composition and geometry, likely act in a compensatory fashion to achieve Rubisco condensation at similar protein concentrations in *Chlamydomonas* and *Chlorella*. Other studies have explored related themes^48^, though direct relationships between binding site geometry on a globular protein and phase separation behavior are scarcely reported. More generally, Rubisco partitioning into the condensed phase depended on both Linker protein concentration and sticker number, as observed previously^49^. Notably however, although we observed uncharacteristic behaviour of the 4x CsLinker variant, we did not observe compelling evidence to support the previously modelled “magic number” effects in our studies^22,50^.

The impact of sticker number, or valency, on phase behaviour has been well-studied in the homotypic phase-separating systems, such as prion-like domains^15,51^ and unfolded protein deposits^52^, where single-residue mutations are often used to modulate valency. By contrast, valency has been less explored in heterotypic systems, where phase separation arises from interactions between distinct molecular species. Nevertheless, several studies demonstrate that valency critically influences condensation in such systems. For example, increasing the number of SH3–PRM binding motifs lowers the critical concentration for LLPS^7^, while RNA-binding valency modulates assembly of RNP granules^53,54^. Similar valency-dependent effects have been modeled theoretically and computationally for heterotypic IDR-IDR mixtures^55^, as well as in a Rubisco-Linker protein context^13,29^. Here, we extend these principles by assessing the effect of valency in the context of other parameters (geometry and affinity) and their impact on biological fitness, which has scarcely been elucidated.

Together, these studies highlight a general principle consistent with our results: increasing the number of interaction motifs enhances network connectivity and lowers the threshold for phase separation. Unlike prior work on carboxysome Linker proteins, where two stickers are sufficient for Rubisco condensation^25,26^, we observe a higher valency requirement in our studied pyrenoid systems, with a minimum of three stickers needed for *in vitro* Rubisco phase separation and four for condensation *in vivo* in *Chlamydomonas*. Given that two sticker fragments bind Rubisco with affinities below their tested concentrations (*i.e.* <10 µM) and substantially higher than single sticker fragments (*K*_D_ >100 µM)^13,27^, it is likely they preferentially bind adjacent subunits on the same Rubisco, as opposed to cross-linking other Rubiscos in the system. Introduction of a third sticker and additional spacer overcomes this threshold—behaviour that was also captured in our LAMMPS simulations. This points towards unexplored effects of sticker geometry on the Linker protein but resolving the biophysical mechanism of this behaviour will require future study.

### Cryo-ET reveals the molecular grammar of pyrenoid organization

Using *in situ* cryo-ET, we correlated the sticker number of EPYC1 variants to the packing density of Rubisco in the pyrenoid. Accurately detecting subtle but significant differences in Rubisco packing between 4x, 5x, and 9x lines required highly accurate particle identification, minimizing false positives (incorrectly assigned Rubisco) and false negatives (missed Rubisco)-an ongoing challenge at the frontier of cryo-ET analysis, especially for biomolecular condensates such as pyrenoids, stress granules^56^, chromatin^33,57–59^, and autophagic condensates^35^. By combining advances in high-resolution template matching^60–62^ with precise masking of the pyrenoid matrix, we achieved high-confidence Rubisco identification, as evidenced by the separation of false positives in our cross-correlation scoring (Fig. S13), and the 13 Å resolution of our subtomogram Rubisco average (Fig. S13b). Interestingly, recent cryo-ET studies of *Chlamydomonas* pyrenoids have been able to push the resolution of Rubisco subtomogram averaging to 7–8 Å resolution, but only by excluding the majority of particles through under-picking or extensive classification^39,40,63^. This reflects the challenge of averaging small particles *in situ*, but also structural heterogeneity caused by the sub-stoichiometric binding of Linker proteins to different sites on the Rubisco surface. The distribution of Rubisco complexes that we observe in our tomograms, including the ∼13 nm preferred distance between first neighbours in all strains and the increased crowding in strains with higher sticker number, is consistent with a liquid-like linked network^22^, rather than randomly positioned particles. More generally, our quantification of Rubisco concentration in the pyrenoid matrix emphasises the importance of low affinity motifs in the context of condensate assembly.

In the chassis strain lacking the EPYC1 Linker protein, the pyrenoid matrix is greatly reduced under both autotrophic and heterotrophic conditions (Fig. S12). However, it is striking that the Rubisco condensate still persists in a reduced form, together with other characteristic structural features of the pyrenoid, including the pyrenoid tubules (and internal minitubules) that intersect the matrix, as well as the surrounding starch sheath. Interestingly, the tubules and starch cover a smaller area than in a WT pyrenoid, apparently restricted to the proximity of the reduced Rubisco matrix. A plausible explanation would be that some degree of Rubisco condensation is maintained by other less abundant proteins, such as MITH1, SAGA1/2, and RBMP1/2, which cross-link Rubisco to the membrane tubules and the starch^36,37,64^. In this manner, the assembly and extent of the Rubisco condensate, pyrenoid tubules, and starch sheath may be co-dependent on each other. We also observed ribosomes around the periphery of the pyrenoid, which have previously been implicated in synthesis of the Rubisco large subunit^65–67^. The exclusion of ribosomes is a common hallmark of phase separation observed by cryo-ET^56,68–70^, and this certainly holds true for the WT pyrenoid matrix. However, in the chassis strain, we observed intermixing of Rubisco and ribosomes, demonstrating how the lack of EPYC1 crosslinking may loosen the residual Rubisco condensate and allow inclusion of Ribosomes.

### Rubisco dynamics and the link to carbon fixing potential

The dynamic nature of Rubisco within the pyrenoid is well established and has been suggested to be essential for its function^22^. Here, we applied single molecule tracking to reveal differences in Rubisco diffusivity that were otherwise inaccessible by ensemble FRAP. Interestingly, in contrast to previous reports, where increased valency is typically associated with reduced diffusivity within condensates^7,71–73^, we observed faster diffusivity of Rubisco in condensates formed with a higher valency Linker protein, both *in vitro* and *in vivo* (Fig. 3e,f). Given that these measurements were performed at fixed Linker protein:Rubisco ratios, the total number of stickers varies. One plausible explanation is that, for 5x and 9x Linker proteins, the finite number of Rubisco binding sites (eight per molecule) is exceeded by the available Linker protein stickers in the system, leading to increased competition for binding sites without increasing cross-linking. Notably, this observed increase in mobility was not reflected in our LAMMPS simulations (Fig. S15), likely because they do not enforce steric or occupancy constraints. Despite this, the lack of discernible carbon fixation efficiency and growth phenotype between the 4x and 5x lines indicates that Rubisco mobility within the condensate is not a critical determinant of ‘steady-state’ Rubisco condensate function, as evidenced by the ordered arrangement of Rubisco in carboxysomes^74–76^, and the apparent lack of mobility of Rubisco in shell-encased diatom pyrenoids^77^. However, this may have implications over division cycles, where Rubisco mobility is essential for pyrenoid inheritance to daughter cells in *Chlamydomonas*^22,42^.

To this end, our study uncovers the physiological relevance of sticker valency in the temporal dynamics of condensate formation, with a direct link to condensate function. Time-resolved assays demonstrate the importance of a fifth sticker, suggesting that its presence accelerates both Rubisco condensation *in vitro* and recruitment into the pyrenoid *in vivo*, with a direct effect on the efficiency of CO_2_ fixation (Fig. 4e). Linker proteins with more than five stickers (e.g., 9x) further increase the rate of recruitment, suggesting that valency can modulate the kinetics of condensate assembly. Although the timescales of our observations *in vitro* and *in vivo* are clearly disconnected (minutes vs hours), we reason that these findings may be unified by recent work showing kinase-mediated dissolution of the pyrenoid in *Chlamydomonas*^42^. Given that 4x represents an apparent threshold for condensation *in vivo* (Fig. 2b), any residual phosphorylation that impacts the effective valency of the Linker protein will affect this variant upon the cellular signal to induce Rubisco condensation. Under steady-state conditions, 4x EPYC1 is sufficient for Rubisco recruitment, but the fifth sticker likely provides robustness against partial Linker protein phosphorylation during early pyrenoid assembly, conferring a potential fitness advantage under fluctuating light and CO_2_ conditions. This observation also aligns with patterns in prokaryotic carboxysomes, whose Linker proteins typically have fewer stickers, consistent with the reduced need for dynamic dissolution in these more static organelles^78,79^.

### A tuneable parameter in the black box of evolution

Together, our findings support the view that sticker number as a tuneable biophysical parameter within the vast, multidimensional landscape of cellular evolution. The relationships we observe between valency, Rubisco packing, mobility, and condensate growth rate illustrate how small molecular changes can propagate across scales to influence cell physiology and ultimately, fitness. Yet, the lack of a simple monotonic relationship between valency and phenotype, most notably the similar fitness of 4x and 5x variants, suggests that changes in sticker number likely balance multiple, sometimes opposing, constraints.

Our data reveal an interesting paradox: natural EPYC1 has evolved essentially the minimum number of stickers required for robust pyrenoid formation, yet variants with more stickers outperform it under our specific competition assays. Why has higher valency not been selected in nature? While increasing valency can accelerate Rubisco recruitment and may enhance CCM performance under specific conditions, additional stickers beyond the functional threshold may provide diminishing or context-dependent returns. Linker proteins are among the most abundant proteins in the cell (0.3-1% of total cell protein)^27,28^; consequently, producing longer, higher-valency variants imposes a substantial biosynthetic cost. Thus, the benefits of increased valency under certain environmental conditions (as we used in our study) may be outweighed in other conditions where faster assembly or higher packing density does not confer a proportional advantage. Our results also raise the possibility that elevated valency could influence the reverse process of pyrenoid dissolution. We observe a clear valency-dependent acceleration of pyrenoid matrix condensation, suggesting that the corresponding dissolution dynamics may also be altered. Given the known kinase-dependent disassembly of the pyrenoid during cell division and environmental transitions, understanding how valency affects disassembly kinetics will be important for future work.

More broadly, our experiments were performed under controlled conditions in which carbon availability was the primary selective pressure. In the natural environment, algal cells inhabit a far richer evolutionary landscape defined by fluctuating CO_2_, light, temperature, nutrient availability, pathogens, and the rates at which these conditions change. These variables, together with regulatory mechanisms such as EPYC1 phosphorylation and differential expression, provide additional levers for tuning condensate behaviour without altering sequence. Mapping how these pressures jointly shape condensate properties will be essential for understanding the evolutionary trajectories available to pyrenoid-forming algae.

This tuneability is particularly relevant when we consider the slow evolutionary rate of Rubisco’s catalytic properties^80^ compared to the rapid evolution of intrinsically disordered proteins, which frequently diversify through repeat expansion^81,82^. Modifying valency through repeat gain or loss represents a relatively simple evolutionary strategy for adjusting condensate assembly, dynamics, or robustness, especially once a binding interface to Rubisco has been established. The high similarity among repeats in Linker proteins is consistent with such a mechanism.

The pyrenoid therefore provides a uniquely tractable system in which to connect the molecular physics of multivalency and phase separation with cellular physiology and evolutionary logic. The same principles, where valency, geometry, and affinity jointly determine condensate function, likely extend to many other biomolecular condensates, from the centrosome to the nucleolus. Ultimately, evolution may not optimise any single condensate property but instead navigate a landscape shaped by competing biochemical and physiological constraints. Sticker number, as shown here, represents one such axis of variation—a molecular tuning parameter through which cells balance stability, dynamics, and cost. Understanding how such parameters emerge, are regulated, and are co-opted across biological systems will help reveal the general principles governing biomolecular condensation in living cells, especially where dynamic assembly and disassembly is essential.

## Materials and Methods

### Linker protein sticker number analysis

Following ref.^83^, all CcmM sequences from β-carboxysome operons (those containing CcmK, CcmL, CcmM and CcmN) were downloaded from all unclassified finished and permanent draft genomes in the Integrated Microbial Genomes and Microbiomes database^84^ on July 16, 2024 (817 genes). Duplicate sequences were removed and annotated Rubisco_small domains (pfam00101) were counted in the remaining 594 genes. Annotated CsoS2 sequences (pfam12288) from α-carboxysome operons (containing CsoSCA, shell hexamers and pentamers) were downloaded on July 17, 2024 (811 genes). Sequences were manually curated to exclude sequences with truncated N-terminal domains, and 35 misannotated open reading frames were manually corrected. After deduplication, the remaining 669 sequences were aligned using MAFFT 7.430 (BLOSUM62 scoring, 1.53 gap penalty, 0.123 offset) in Geneious Prime 2024. N-terminal domains were extracted and used for MEME searches to determine the number of Rubisco-binding regions using the following parameters (-protein - mod anr -nmotifs 1 -minw 6 -maxw 20 -objfun classic -markov_order 0 -csites 6000 -V -p 11). Homologs of EPYC1 and CsLinker were obtained by BLASTp searches against the NCBI non-redundant sequences (nr) database, and the JGI Phycocosm database. Duplicate sequences were removed and the number of stickers for each species was counted.

### Design of Linker protein variants

To design the EPYC1 and CsLinker variants we removed the chloroplast transit peptide and centred each repeat on the corresponding Rubisco-binding helix such that a full repeat spanned half of the preceding spacer and half of the following spacer. Because the C-terminal repeats of both EPYC1 and CsLinker are less conserved, these were removed first when generating shorter variants. Conversely, for extended variants, additional repeats were appended at the N-terminus. In these cases, we selected the most representative repeat sequences and preserved consecutive repeat order wherever possible.

### Cloning and purification of Linker protein variants

Untagged EPYC1 and CsLinker variants were codon-optimised for *E. coli* and synthesised by TWIST Bioscience either directly in the expression vector or as gene fragments, which were subsequently cloned into the same N-terminal mEGFP-TEV, C-terminal TEV-MBP fusion expression vector by Golden Gate cloning with BsaI. BL21(DE3) cells carrying expression plasmids were cultured in Luria Broth at 37 °C to an OD_600_ of 0.5-0.8 before induction with 1 mM IPTG either at 37 °C for 2-4 hours or overnight at room temperature. Cells were lysed by ultrasonication in lysis buffer (50 mM Tris-HCl, 500 mM NaCl, and 25 mM imidazole at pH 8.0) and the soluble fraction and applied to 5 mL HisTrap™ columns (Cytiva Life Sciences) equilibrated with lysis buffer at 4 °C. Proteins were eluted using a gradient to 100% elution buffer (50 mM Tris-HCl, 500 mM NaCl, 500 mM imidazole at pH 8.0) over 20 column volumes. The mEGFP and MBP fusions were cleaved by overnight incubation with purified TEV protease^85^ at ∼1:100 w/w ratio on ice. Cleaved samples were buffer exchanged into lysis buffer by concentration using Amicon Ultra-15 10 kDA MWCO centrifugal filter units (Merck) and applied to HisTrap columns. The flowthrough fractions were collected and subjected to size-exclusion chromatography (SEC) on a HiLoad 16/600 Superdex 75 prep grade column (Cytiva Life Science) equilibrated in SEC buffer (50 mM Tris-HCl, 500 mM NaCl, 5% glycerol (w/v)). Samples were stored at −70 °C until use. Protein concentrations were determined by Bradford assay.

### Purification of Rubisco

*Chlamydomonas reinhardtii* (CC-4533) and *Chlorella sorokiniana* (SAG 211-8k) Tris-acetate-phosphate (TAP) medium with revised trace elements^86^ sparged with 3% CO_2_ to mid-exponential phase (2-8×10^6^ cells mL^-1^). Cells were harvested by centrifugation and resuspended in Rubisco lysis buffer (50 mM Tris-HCl, 100 mM NaCl, 0.5 mM EDTA at pH 8.0) supplemented with 5 mM DTT and 2 mM PMSF. Cells were lysed by ultrasonication ( *C. reinhardtii*) or cell disruptor (*C. sorokiniana*). The soluble fraction after centrifugation (30,000 *g* for 30 min) was subjected to two sequential ammonium sulfate precipitations: first at 190 mg mL^-1^, then at 390 mg mL^-1^, each for 1–2 h at 4 °C. The pellet from the second precipitation was resuspended in ion-exchange (IEX) buffer A (50 mM Tris-HCl, 50 mM NaCl, 1 mM DTT at pH 8.0) and loaded onto 10%-30% sucrose (w/v) gradients prepared in the same buffer. Gradients were centrifuged at 37,000 rpm for 16.5 hours at 4 °C (SW 41 Ti rotor, Beckman Coulter). Rubisco-containing fractions were pooled and applied to a 5 mL HiTrap Q XL column (Cytiva Life Sciences) that was pre-equilibrated with IEX buffer A, and proteins were eluted with IEX buffer B (50 mM Tris-HCl, 500 mM NaCl, 1 mM DTT at pH). Rubisco fractions were pooled and concentrated using Amicon Ultra-15 centrifugal filter units (Merck), and subjected to size-exclusion chromatography (SEC) on a HiLoad 16/600 Superdex 200 prep grade column (Cytiva) equilibrated in SEC buffer (50 mM Tris-HCl, 50 mM NaCl, 5% (v/v) glycerol, 1 mM DTT at pH 8.0). Protein concentrations were determined using a NanoDrop, and samples were stored at −70 °C. The same process was completed for the purification of the Rubisco / Rubisco-mVenus pool from the chassis strain.

### Atto594 labelling of Rubisco

Rubisco was buffer-exchanged into labelling buffer (50 mM HEPES-KOH, pH 7.4, 50 mM NaCl), and 1-10 mg of lyophilized Atto 594 NHS-ester dye (Merck, 68616-1KT) was resuspended directly into the protein solution using a 200 µL pipette tip. Labelling was carried out at room temperature for 2 hours with gentle agitation. The reaction mixture was applied to a Superdex 200 Increase 10/300 GL column (Cytiva) equilibrated with SEC buffer (50 mM Tris-HCl, 50 mM NaCl, 5% glycerol (w/v), 1 mM DTT at pH 8.0) to separate labelled Rubisco from unreacted dye.

### Fluorescence microscopy for visualization of phase separation

Sticker variants were buffer-exchanged into condensate buffer (50 mM Tris-HCl, 50 mM NaCl at pH 8.0) immediately before the start of the experiments. Rubisco was prepared at a final concentration of 1 µM (including 5 nM Atto594-labelled) and mixed with varying concentrations of sticker variants in a total reaction volume of 5 µL using condensate buffer. The reaction mixture was immediately drop-cast onto individual wells of an ibidi 15-well µ-Slide. Samples were visualized using an inverted fluorescence microscope (Zeiss Axiovert 200) with a 63x objective under the TRITC channel to detect droplets. Each reaction volume was scanned for droplet formation, and images were captured 5 minutes after mixing.

### Droplet sedimentation assays

Sticker variants were buffer-exchanged into condensate buffer (50 mM Tris-HCl, pH 8.0, 50 mM NaCl) immediately prior to the assay. Rubisco (1 µM final concentration) was mixed with varying concentrations of sticker variants in a 5 µL reaction volume using the same buffer. Samples were incubated at room temperature for 20 minutes to allow condensates to form, then centrifuged at 18,000 *g* for 20 minutes at RT. The supernatant (dilute phase) was carefully removed, and the pellet was resuspended in a 1xSDS loading buffer. Both fractions were analyzed by SDS-PAGE, and band densitometry was performed using Fiji. For time-resolved sedimentation assays, droplet mixtures were centrifuged at 30,000 *g* for 1 minute following the indicated incubation period.

### Atomic force microscopy (AFM)

Atomic force spectroscopy of EPYC1 was carried out using an EPYC1 construct which was modified by Q5 site-directed mutagenesis (NEB) to sequentially add an N-terminal 6xHis tag and a C-terminal cysteine residue. The construct was purified as for the other EPYC1 variants, except a cation exchange step was performed using a 5 mL Capto S column. The elution from the nickel affinity purification was diluted 10-fold in CX buffer (50 mM Na-phosphate pH 7.5), bound to the column and washed with CX buffer C (50 mM Na-phosphate pH 7.5, 20 mM NaCl) and eluted with a gradient to CX elution buffer (50 mM Na-phosphate pH 7.5, 1M NaCl) over 10 CVs. For biotinylation, the sample was diluted to 1 mg mL^-1^ in PBS supplemented with 1 mM DTT and incubated at 4 °C for 2 h, before precipitation with ammonium sulphate (70% saturation, 10 m mixing, centrifuged at 13,000 *g* for 10 m at 4 °C). The pellet was washed with 1 mL of wash buffer (100 mM NaPO_4_ pH 7.3, 200 mM NaCl, 1 mM EDTA, 70% ammonium sulphate), centrifuged, and resuspended in 2.5 mL of wash buffer containing 5x molar excess of Biotin-dPEG-Mal (Sigma QBD10201) and incubated at RT for 30 m. The reaction was quenched with β-mercaptoethanol (0.5% v/v) for 10 min at RT. Free biotin was removed by passing the sample over a 1 mL HisTrap column. The eluted protein was buffer exchanged into storage buffer (50 mM Tris HCl pH8, 500 mM NaCl, 5% glycerol), and the efficiency was measured using Strep-Tactin resin (70% efficient).

Experiments were performed using a JPK NanoWizard II AFM equipped with NPG-10 probes (cantilever D, Bruker). Probes were UV-ozone cleaned for 30 min, then functionalized overnight with a self-assembling monolayer of OEG-thiol:biotin-OEG-thiol (1 mM in ethanol, 98:2 molar ratio)^87^. Silicon wafers were UV-ozone cleaned for 30 min, mounted to Teflon-coated metal discs, and coated with small unilamellar vesicles (2 g mL^-1^ DOPC:DOGS-NTA, 99:1 or 99.5:0.5 mol:mol, 5 mM NiCl_2_, in HBS buffer [10 mM HEPES pH 7.4, 150 mM NaCl]) for 30 min to form supported lipid bilayers^88^. Bilayers were washed 5x with HBS and 5x with TBS (50 mM Tris pH 8.0, 50 mM NaCl). Functionalized probes were incubated with 40 µg mL^-1^ streptavidin in TBS for 10 min, washed 5x in TBS, then incubated with 20 µg mL^-1^ EPYC1 in TBS for 20 min before a final wash. Force curves were collected at room temperature at approach and retract velocities of 200 or 400 nm/s with no dwell time and a maximum loading force of 300 pN. Hundreds of curves from five independent experiments were analysed using the JPK Data Processing software to extract contour length (L*_c_*) and persistence length (L*_p_*) using a worm-like chain model. Only curves with fitted L*_c_* values within 25% of the expected EPYC1 contour length were retained. A consensus L*_p_* was determined from the peak of bootstrapped probability density functions generated using SciPy’s gaussian_kde.

### Large-scale Atomic/Molecular Massively Parallel Simulator (LAMMPS) simulations

Simulations were performed with the 2Aug2023 LAMMPS release on the Viking supercomputer. Rubiscos were modelled as 13 nm rigid body spheres with binding sites of 1 nm radius arranged according to their experimentally determined positions^23,27^ that together could rotate and diffuse freely. EPYC1 and CsLinker consisted of chains of sticker spheres separated by spacer spheres, all connected with FENE bonds, following^29^. Stickers may interact with Rubisco binding sites at distance *r* < *r*_0_ = 1 nm via a shifted cosine potential U(*r*)= ½ *A* [1+cos(*π r*/*r*_c_)], where *A* is the well depth. In our model, sticker and spacer beads do not participate in excluded-volume interactions with each other, so that the polymer is effectively a freely-jointed chain. All other excluded volumes are as in GrandPre et al. Spacer beads participate identically to sticker beads except they cannot interact with Rubisco binding sites.

Sticker interactions were parameterised against SPR-derived *K*_D_ values by placing a single Rubisco and a single sticker bead in simulation boxes of varying size to change the effective concentration, over which the potential depth was also changed. The proportional interaction time was used to calculate the *K*_D_, and the interaction well depth corresponding to the experimental *K*_D_ was used (*Chlorella*: −58.96 pN nm; *Chlamydomonas*: −53.68 pN nm; effective well depths 14.2 and 13.0 kT, respectively) (Fig. S7a). The FENE bond spring constants were tuned against SPR data of 2 sticker fragments binding to Rubisco, where the maximum permitted extension was 20 nm (Fig. S7b).

For full simulations, boxes of 130×130×210 nm or 85×85×210 nm into which either 832 or 325 Rubisco molecules respectively were inserted on a lattice were used. Linker polymers were introduced at chosen Rubisco:Linker ratios and the system was relaxed using conjugate gradient minimisation to remove overlaps, before simulation of 50,000,000 timesteps (d_t_ = 0.05 ns) with periodic boundary conditions, and with the interaction potential strength linearly increasing across the simulation between 0 and its target well depth. This in effect “annealed” the molecules in a condensed phase. The simulation box was then expanded to either 130×130×500 nm or 85×85×500 nm to give the proto-condensate room to dissolve, and run for 50,000,000 more simulation steps. Finally, production runs of >30,000,000 simulation steps were performed. In both equilibration and simulation steps, snapshots were written every 100,000 steps. All simulations were carried out at 300 K, maintained with a Langevin thermostat with damping parameter 5 as in GrandPre et al.

Analysis was performed using bespoke Python scripts which constructed the full periodic system box and calculated for each Rubisco the number of other Rubiscos closer than 20 nm, following^22^, confirmed by radial distribution function analysis (Fig. S7d). Given the large edge effect of the condensate in the simulation box, the threshold for the number of neighbouring Rubiscos within 20 nm determined to be within the condensate was determined analysing neighbour count distributions which showed a clear delineation between dilute and condensed Rubiscos (Fig. S7c). A threshold of 5 neighbours was used in all subsequent analyses. For diffusion measurements, diffusivity rates in the condensed and dilute phase were separated by fitting two gaussians to the distributions of diffusivity rates, in which the faster diffusing population was assumed to be in the dilute phase. Single-molecule diffusion coefficients were calculated from the mean squared displacement (MSD) of Rubisco molecules. For each molecule, coordinates from the last portion of the trajectory were used. MSDs were computed for multiple lag times by averaging squared displacements between all frame pairs separated by the lag. The linear portion of the MSD versus time curve was fitted to extract the diffusion coefficient D using MSD = 6Dt in three dimensions. Visualisation of the simulations were completed using Ovito.

### Statistical physics models

In our LAMMPS simulations, the phase behaviour of Rubisco:Linker systems is controlled by the sticker-site interaction, spacer flexibility, and the concentrations of both constituents. As mapping out the full phase diagram is computationally infeasible, we used the Linker protein-mediated pairwise Rubisco-Rubisco interaction as a numerically tractable proxy, represented by full concentration-dependent dimerization diagrams (see ref.^13^). The model is parametrised using the dissociation constant *K*_D_, the sticker-site binding energy ε, and the Kuhn length of amino acids, *l*_K_ (these can be mapped onto the parameters used in LAMMPS, see Supplementary Note 1). In addition to these properties the spatial coordinates of sites (and corresponding intersite distances) are parametrised using a geometric model for Rubisco.

The dissociation constant was parametrised using the titration data of single-sticker constructs (*Chlamydomonas*: 414 μM; *Chlorella*: 130 μM). The intersite distances were generated using a ‘cube model’ (for which the sites are at the corner of a cube at distance 6.7 nm from the centre of mass; this model previously provided encouraging predictions for the critical concentration of phase separation), and a ‘sphere model’ for which the sites are at the same locations as in the LAMMPS simulations (see Supplementary note and Fig. S8). The remaining two parameters (ε and *l*_K_) were determined using the titration of multi-sticker constructs. For *Chlamydomonas* this yielded *l*_K_ = 0.88 ± 0.02 nm and ε = −11.8 ± 0.1 *k*_B_*T* for the cube (as in ref.^13^) and *l*_K_ = 0.58 ± 0.01 nm and ε = −12.5 ± 0.1 *k*_B_*T* for the sphere. For *Chlorella* no unique curve fit could be obtained; for this system ε = ε0 *l*_K_^α^ with ε0 = −11.97 ± 0.02 *k*_B_*T* and α = 0.3543 ± 0.004 led to a continuous series of parametrisations (in the physically relevant range of *l*_K_ from 0.5 to 1 nm, the corresponding binding energy ranged from −15.2 to −11.9 *k*_B_*T*).

In Fig. S8 we find the choice of geometry (cube vs sphere) affects the prediction for phase separation (discrepancies suggest a role excluded-volume interaction, non-specific interactions, and/or multi-Rubisco interactions leading to (anti-)cooperative self-assembly). More importantly, we find the dependence of the critical concentration versus the number of stickers is robust against variations in the geometry (cube vs sphere), as well as a robustness of the results for variations of *l*_K_ and ε along the ε = ε_0_ *l*_K_^α^ parametrisation.

### Chassis creation

An RbcS2-mVenus expression construct was made by Golden Gate assembly of the RbcS2 (Cre02.g120150) sequence with the hybrid HSP70/RbcS2 promoter (pAR, pCM0-015) and RbcS2 terminator (pCM0-115)^89^ to create a level 1 expression part. This vector was assembled with a hygromycin resistance cassette into pAGM4723 at level 2. The chassis was created by nuclear transformation of the *epyc1* knockout (CC-5360) with this construct, using electroporation following^90^, and selection on TAP + hygromycin (25 µg mL^-1^) plates.

### Cloning and chloroplast transformation of EPYC1 sticker variants

EPYC1 sticker variants were codon-optimised using Geneious Prime using the *Chlamydomonas* chloroplast codon usage table (Kazusa). Variants were synthesised by TWIST Bioscience and Golden Gate assembled into pME_Cp_2_098^30^. Some shorter variants (6x, 7x, 8x) were made by PCR amplification of the 9x sequence and reassembly into the expression vector. Chloroplast transformations were performed by particle bombardment with a Biolistic PDS-1000/He system (Bio-Rad). For each bombardment, 0.5 mg of 550 nm gold nanoparticles (Seashell Technologies) were coated with 1 µg of plasmid DNA, following the manufacturer’s protocol. Freshly grown cells in their exponential phase (∼0.5-1×10^7^) were plated on circular TAP plates containing 150 µg mL^-1^ of either spectinomycin or tobramycin and positioned ∼9 cm below a 1,100-psi rupture disk. Selection was performed under medium light intensity (∼50 µmol photons m^-2^ s^-1^) until colonies appeared, after which 12 colonies per transformation were restreaked to homoplasmy on TAP agar plates supplemented with 250 µg mL^-1^ of either spectinomycin or tobramycin. Homoplasmic integration of all the variants was confirmed by colony PCR (Fig. S10b).

### Rubisco partitioning measurement

Cells were grown in TAP medium in 25 mL flasks to mid exponential phase before centrifugation and resuspension in TP minimal medium for different time points prior to imaging. 10 µL of cells were immobilised in ibidi µ-Slide 18-well chambered coverslips using 40 µl of 1% (w/v) TP low-melting-point agarose. Images were captured using a Zeiss LSM 980 confocal microscope in Airyscan mode with a 63x oil immersion lens. Images were in Fiji and Rubisco partitioning was calculated as described previously, with the addition of a correction for background chloroplast fluorescence (Fig. S11b)^27^.

### Cryo-ET sample preparation and data acquisition

A total of 4.5 µL cell suspension (1500–2000 cells/µL), grown in TP minimal media at air level CO_2_ and 100 µE light, were applied to a holey carbon R2/1 grid (Quantifoil). Cells were plunge-frozen in liquid ethane with back-sided blotting using a Leica EM GP2 (2 s blot time and 90% humidity). Samples were stored under liquid nitrogen conditions.

FIB milling was performed as described in^91^. Briefly, grids clipped in Autogrids (Thermo Fisher Scientific) were loaded into an Aquilos 2 microscope (Thermo Fisher Scientific) and a layer of organometallic platinum was applied using the gas injection system. For milling and imaging, the gallium ion beam was operated at 30 kV.

Tilt-series were acquired on a Titan Krios G4 transmission electron microscope operating at 300 kV equipped with a Selectris X energy filter with a slit set to 10 eV and a Falcon 4i direct electron detector (Thermo Fisher Scientific), recording dose-fractionated movies in TIFF format. A dose-symmetric tilt scheme was used, set by the TEM Tomography 5 software (Thermo Fisher Scientific) in a tilt span of ± 54°, covered by 2° steps starting at either ± 10° offset to compensate for the lamella pre-tilt. Target defocus was set for each tilt-series in a range of −2 µm to −4 µm in steps of 0.5 µm. The microscope was set to a nominal magnification of 53,000x, corresponding to a pixel size of 2.42 Å at the sample level and a nominal dose of 2 e^-^/Å^2^ per tilt image. A 100 μm objective aperture was inserted during collection.

Tomograms were entirely processed using Scipion^92^ using the workflow as described^93^. Briefly, raw frames were motion corrected using Motioncorr3^94^, and images were aligned with AreTomo2^95^ after manual tilt-series curation. CTF-corrected (by phase-flipping), dose filtered, and four times binned tomograms were reconstructed using IMOD^96^.

Tomograms were denoised from their reconstructed half-volumes using DeepDeWedge^97^. From the denoised tomograms, membrane segmentation was done with Membrain-seg^98^, and precise pyrenoid matrix masks were drawn manually in Amira (Thermo Fisher Scientific) and used for pyrenoid volume calculation. Rubisco particle identification was performed on CTF-corrected tomograms using pytom-match-pick^60^. Template matching was performed with the map published in^99^, scaled to bin4 pixel size and lowpass filtered to bin4 Nyquist frequency. An average produced in RELION5^100^ with the initial TM particle picks (see Supplementary) was used to re-run TM and to confirm that the particle selection was unambiguous. Cross-correlation score based extraction thresholds were calculated using a custom Python script. The final particle coordinates were used for the spatial analysis.

### Cryo-ET data analysis

Rubisco density was calculated as the fraction of Rubisco particle number and volume of the pyrenoid mask, scaled to one cube micron or millilitre. Nearest-neighbor (NN) distances were calculated for each particle of each tomogram using a cKDTree (SciPy)^101^ implementation. The raw NN distance values were extracted and the arithmetic mean as well as the number of neighbours within a given spherical distance was computed as the central measure of the spatial distribution. Scripts for the analysis are available.

### Fluorescence recovery after photobleaching (FRAP)

Cells were prepared for FRAP using the same protocol as for the Rubisco partitioning measurements. Imaging was performed on a Zeiss LSM980 confocal microscope equipped with a 63x objective and the Definite Focus module. Three pre-bleach images were acquired before bleaching approximately 30–50% of the pyrenoid area using two consecutive 500 ms bleach pulses at 15% laser power of the 514 nm line. A total of 250 images were acquired at 1 s intervals to monitor fluorescence recovery. Images were drift-corrected in Fiji using the Image Stabilizer plugin^102^. Background-subtracted mean gray values were extracted from bleached and unbleached regions of interest. Signal homogeneity within a rectangular region spanning both areas was calculated relative to the pre-bleach baseline (Fig. S14). The recovery kinetics were characterized by fitting an exponential model to the post-bleach data, y(*t*) = *A* x (1 - e^-*kt*^), where *A* is the recovery plateau, *k* is the rate constant and *t* is the time after bleaching.

### Single particle tracking

For *in vivo* imaging, cultures of either WT or EPYC1 variant (4x, 5x, 9x) cells in TAP media were transferred to TP minimal media for 48 h under ambient CO_2_, then concentrated (5 min, 1500 *g*) and resuspended. Cells were spotted onto an agar pad on a microscope slide (ThermoFisher GeneFrame) containing TP media + 1.5% low melting point agarose and sealed under a #1.5H precision coverslip.

For in vitro imaging, a 5 µL droplet suspension was produced by mixing recombinant EPYC1 sticker variants (4x, 5x, 9x) with 1 µM of the purified Rubisco / Rubisco-mVenus pool in condensate buffer (50 mM Tris-HCl, pH 8.0, 50 mM NaCl). This suspension was introduced slowly to a tunnel slide^103^ constructed from either plasma-cleaned or Rain-X/Pluronic F-127-passivated coverslips^104^ to prevent complete wetting of condensates.

Cells or droplets sessile on the agar surface were then imaged using SlimVar fluorescence ^41^. For each acquisition, either the pyrenoid of an individual cell, or a single pyrenoid-like droplet was centred in the field of view and illuminated with a 514 nm wavelength laser at high irradiance (kW cm^-2^) and at an oblique angle, in a focal plane >1 µm above the coverslip surface to minimise background. A rapid, brief sequence of 200-5,000 frames was captured, each of ∼5 ms exposure separated by ∼1 ms readout time. This process of intense photobleaching coupled with sufficiently rapid observation of the fluorescent signal made it possible to identify discrete particles (as small as a single molecule of Rubisco-mVenus) within the condensate, to quantify their molecular content and to track their diffusion with minimal motion blur.

Image sequences were post-processed using ADEMScode software^105^ to identify tracks from maxima in fluorescence intensity linked across multiple frames, then to quantify their number of Rubisco-mVenus molecules and their individual diffusivities over time using a mean-squared displacement analysis^41,106,107^.

### Oxygen evolution

For each measurement, a culture volume containing 100 µg of chlorophyll was pelleted at 1,500 *g* for 5 min at 25 °C then resuspended in 2 mL of TP medium that had been depleted of CO_2_ by aeration with CO_2_-scrubbed air. The resuspended culture was transferred to the measurement chamber, which was equilibrated to 25 °C, stirred at 75%, and illuminated at 150 µmol photons m^-2^ s^-1^. The sample was bubbled continuously with CO_2_-scrubbed air for ∼10 min until the rate of oxygen evolution approached zero. Prior to the measurement series, the culture was illuminated at 750 µmol photons m^-2^ s^-1^ until the oxygen signal stabilized again. For each inorganic carbon addition, 2 µL of each solution was injected using a 10 µL Hamilton syringe. After each injection, the syringe was left in the injection port for 30 s to maintain chamber sealing. Injections were performed at 60 s intervals. Data analysis was conducted in a blinded manner using a custom Google Colab notebook. For each injection response, the maximum oxygen evolution rate over a window of at least 10 s post-injection was extracted. Replicates were collected in alternating fashion between samples.

### Spot test assays

Cultures were grown in TAP to mid-exponential phase before harvesting, washing with TP thrice and then serial dilution to ∼100,000, 1,000, and 100 cells mL^-1^ TP. 10 µL of each dilution was spotted onto TP, pH 7.4 minimal agar plates. Plates were incubated under the indicated condition at ∼28 °C for 7 days in chambers maintained at 0.01%, ambient (∼0.04%), or 3% CO_2_. Growth was monitored by imaging plates at the end of the incubation period.

### Liquid growth assays

For all growth phenotyping experiments, TAP-grown, mid-exponential cells were inoculated to 5×10^5^ cells mL^-1^ in TP, pH 7.4 equilibrated to ambient CO_2_ (∼420 ppm). Growth phenotyping of all 12 strains was completed in 12-well plates containing 2.5 mL of TP media, shaken at 190 rpm. Plates were incubated in custom growth chambers maintained at ∼420 ppm by constant supply of compressed air. Optical density at 750 nm (OD_750_) was measured every 24 hours using a SPECTROstar Nano plate reader. Cultures were kept in suspension by constant agitation prior to measurement. The measurement program involved 5 seconds of orbital shaking at 100 rpm followed by averaged spiral measurement of OD_750_ in each well within a diameter of 1 cm at the centre of the well. Triplicate cultures were randomly arranged across 4 12-well plates, which also contained 3 blank wells per plate.

For cell count measurements of 4x, 5x, chassis and WT strains cultures were grown in either 25 mL of TP media, in open 50 mL Erlenmeyer flasks, or 50 mL of media in cotton-plugged 100 mL Erlenmeyer flasks for the 400 ppm and 100 ppm growth assays respectively. Cultures were shaken at 180 rpm in custom growth chambers supplied with constant supply of air at the indicated CO2 concentration, with a replacement rate of 1 chamber volume ∼20 minutes. Triplicate cultures were measured at 24 hour intervals by sampling of ∼100 µL volumes that were counted twice as technical replicates using a Countess 3 automated cell counter.

### Competition growth assays

To create the chloroplast marker lines, mCherry (pLM1507) and mVenus (pME_Cp_0_4_009) fluorophore parts were separately assembled into pME_Cp_1_475 by Golden Gate assembly with BsaI. The resulting vectors were assembled with a Tobramycin resistance cassette (pME_Cp_1_472) and a filler sequence (ATGACGTCTCGAAGCGTCACAAGTACGAGACGGATC) into the acceptor vector pME_Cp_0_7-8_005 using BsmBI. The resulting plasmids were transformed into 4x, 5x, and 9x lines created in the *epyc1* background as described for the chassis. Following transformation colonies were selected on TAP + spectinomycin 250 µg mL^-1^ + tobramycin 250 µg mL^-1^ plates and restreaked to homoplasmy. For the competition assays, cells were grown to ∼1×10^7^ cells mL^-1^ in TAP and relevant strains were mixed in equal volumes and diluted 10-fold in TP media to a final volume of 5 mL in 6-well plates with a concentration of ∼1×10^6^ cells mL^-1^. A baseline measurement was taken by flow cytometry to determine the population distribution of each of the strains in each experiment. The plates were incubated under ambient CO_2_ conditions at ∼100 µmol photons m^-2^ s^-1^. mVenus and mCherry expression were analysed by flow cytometry (CytoFLEX LX, Beckman Coulter). Cells were distinguished from debris by forward (FSC) and side (SSC) scatter signals generated by the 488 nm laser, and singlets were gated using FSC-height versus FSC-area plots. Chlorophyll autofluorescence (488 nm excitation, 690/50 filter) was used to verify intact algal cells. mVenus and mCherry fluorescence were excited with 488 nm and 561 nm lasers, respectively, and detected with 525/40 and 610/20 filters. Data was processed with CytExpert software (Beckman Coulter).

## Acknowledgements

The University of York Department of Biology Technology Facility are acknowledged for providing access/support for fluorescence microscopy and flow cytometry. Members of the Mackinder lab, the YP3 retreat and the CAPP community are thanked for fruitful discussions. G.K., J.B., A.P-D., J.S., C.S., J.H., C.D., M.H., M.J.P., M.C.L. and L.C.M.M. were funded by EPSRC grant EP/W024063/1 as part of the York Physics of Pyrenoids Project (YP3). M.C.L. from EPSRC grant EP/Y000501/1. J.B. from CTRF grant AP23-1_023. O.N. and L.C.M.M. were funded by R/T020679/1. P.V.d.S, M.H. and B.D.E. were funded by Swiss Nanoscience Institute PhD School grant P2204. Additional funding to B.D.E. from ERC consolidator grant “cryOcean” (fulfilled by the Swiss State Secretariat for Education, Research, and Innovation, M822.00045). L.F. was funded 50:50 by the University of York School of PET and EPSRC DTP.

## Data and Software Accessibility

All data associated with the study will be made available by public repositories. cryo-ET subtomogram averages, and cellular tomograms will be made available in the Electron Microscopy Data Bank. All other associated data will be made available in a Zenodo repository. Software is available through separate GitHub repositories: cryo-ET analysis: github.com/Phaips/TMetric

LAMMPS analysis: github.com/james-r-barrett/RubiCon

Oxygen evolution: github.com/james-r-barrett/oxysolve

statistical physics modelling: https://github.com/CharleySchaefer/RubiscoMonomers

SlimVar tracking: github.com/alex-paynedwyer/single-molecule-tools-alpd

## Author contributions

Conceptualization, G.K., J.B., P.V.d.S., C.S., M.J.P., M.C.L., M.H., B.D.E., L.C.M.M.; Formal analysis, G.K., J.B., P.V.d.S., O.N., J.S., A. P-D., C.S., L.F., J.H., V.M.; Funding acquisition, C.S., R.P.R., M.J.P, M.C.L, M.H., B.D.E., L.C.M.M.; Investigation, G.K., J.B., P.V.d.S., O.N., J.S., A. P-D., C.S., L.F., J.H., V.M., R.P.R.; Methodology, G.K., J.B., P.V.d.S., O.N., J.S., A. P-D., C.S., L.F., J.H., V.M., R.P.R.; Software, J.B., P.V.d.S., J.S., A.P-D., C.S., L.F., V.M.; Supervision, J.B., C.S., R.P.R., M.J.P., M.C.L., M.H., B.D.E., L.C.M.M.; Validation, G.K., J.B., P.V.d.S., O.N., J.S., A. P-D., C.S., L.F., J.H., V.M., R.P.R.; Visualisation, G.K., J.B., P.V.d.S., O.N., J.S., C.S., J.H.; Writing – original draft, G.K., J.B., P.V.d.S.; Writing – review & editing, G.K., J.B., P.V.d.S., C.S., M.J.P., M.C.L., M.H., B.D.E., L.C.M.M.;

## Supporting Figures and Tables

**Fig. S1:**
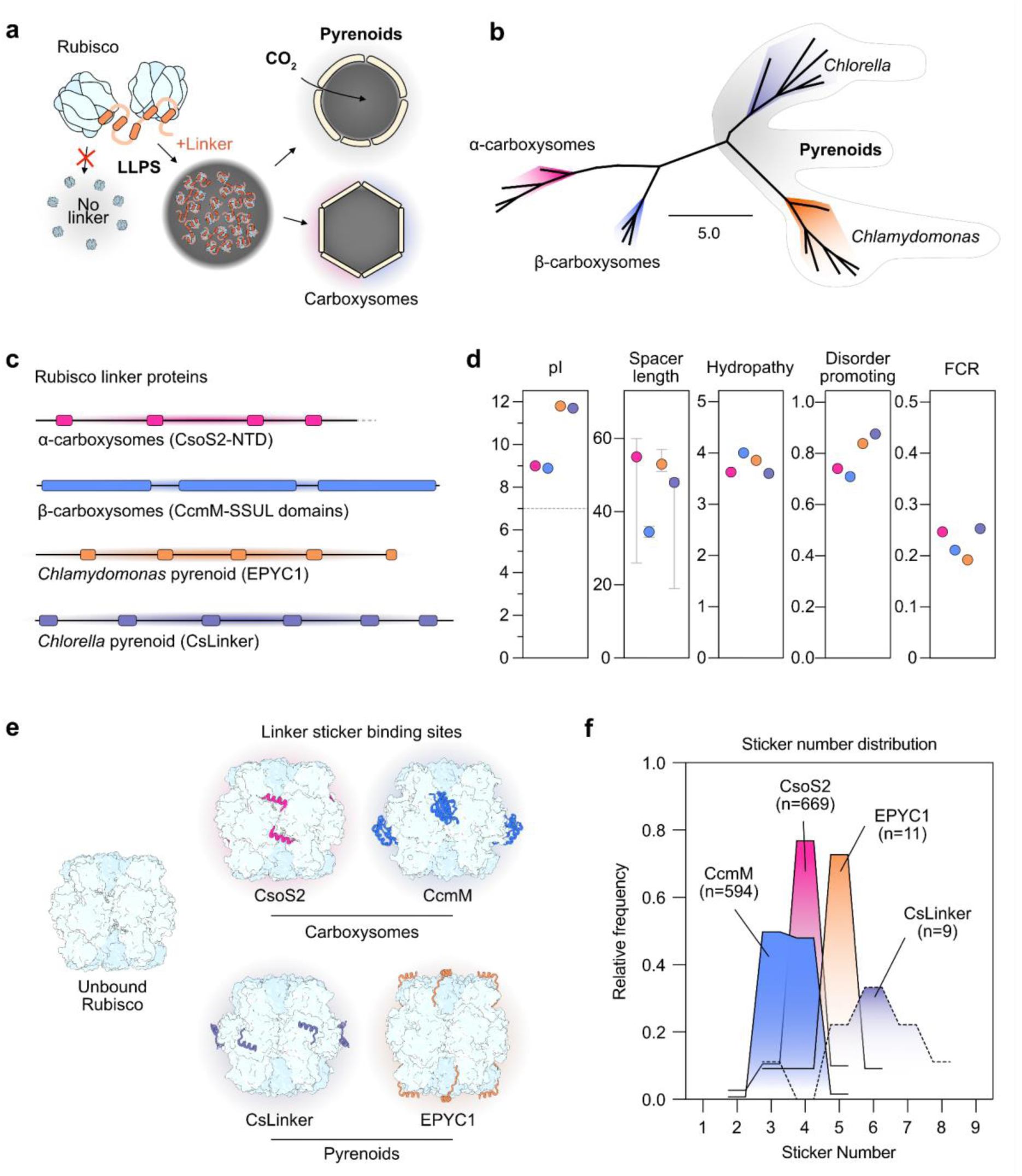
Characteristics of Rubisco-condensing Linker proteins from carboxysomes and pyrenoids. **a)** Linker proteins drive Rubisco condensation by liquid-liquid phase separation (LLPS), forming the pyrenoid matrix and carboxysome condensate. **b)** Phylogeny of Rubisco large subunit sequences from representative carboxysome and pyrenoid species characterised Linker proteins. A consensus tree topology from 1,000 RAxML bootstrap replicates (GAMMA BLOSUM62 model) is shown. **c)** Schematic of Linker proteins, with rectangles indicating Rubisco-binding regions (as shown in panel e). For carboxysome proteins, only the Rubisco-binding regions are depicted. **d)** Physicochemical properties of Linker proteins. Hydropathy, disorder promoting and fraction of charged residues (FCR) were calculated using CIDER^108^. Error bars for spacer length indicate the range among sequences shown in panel c. **e)** Experimentally determined structures of Rubisco-binding regions for each Linker protein bound to their cognate Rubisco. PDB IDs: 6UEW (CsoS2), 6HBC (CcmM), 8Q05 (CsLinker), 7JFO (EPYC1). **f)** Relative frequency of sticker numbers among homologues of each Linker protein.

**Fig. S2:**
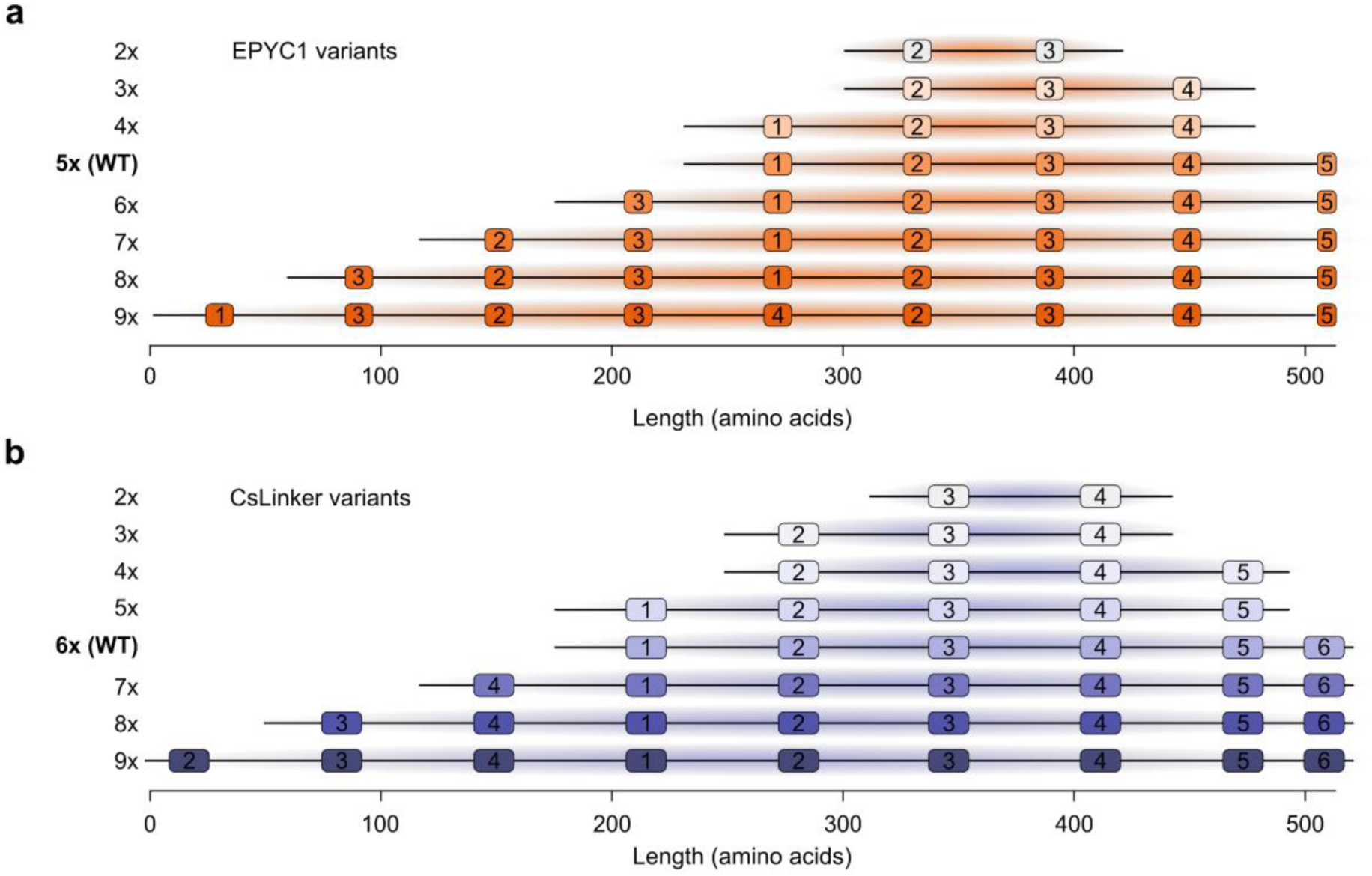
Schematic of sticker variants in EPYC1 (a) and CsLinker (b). Rectangles indicate stickers, numbered according to their position in the WT sequences.

**Fig. S3:**
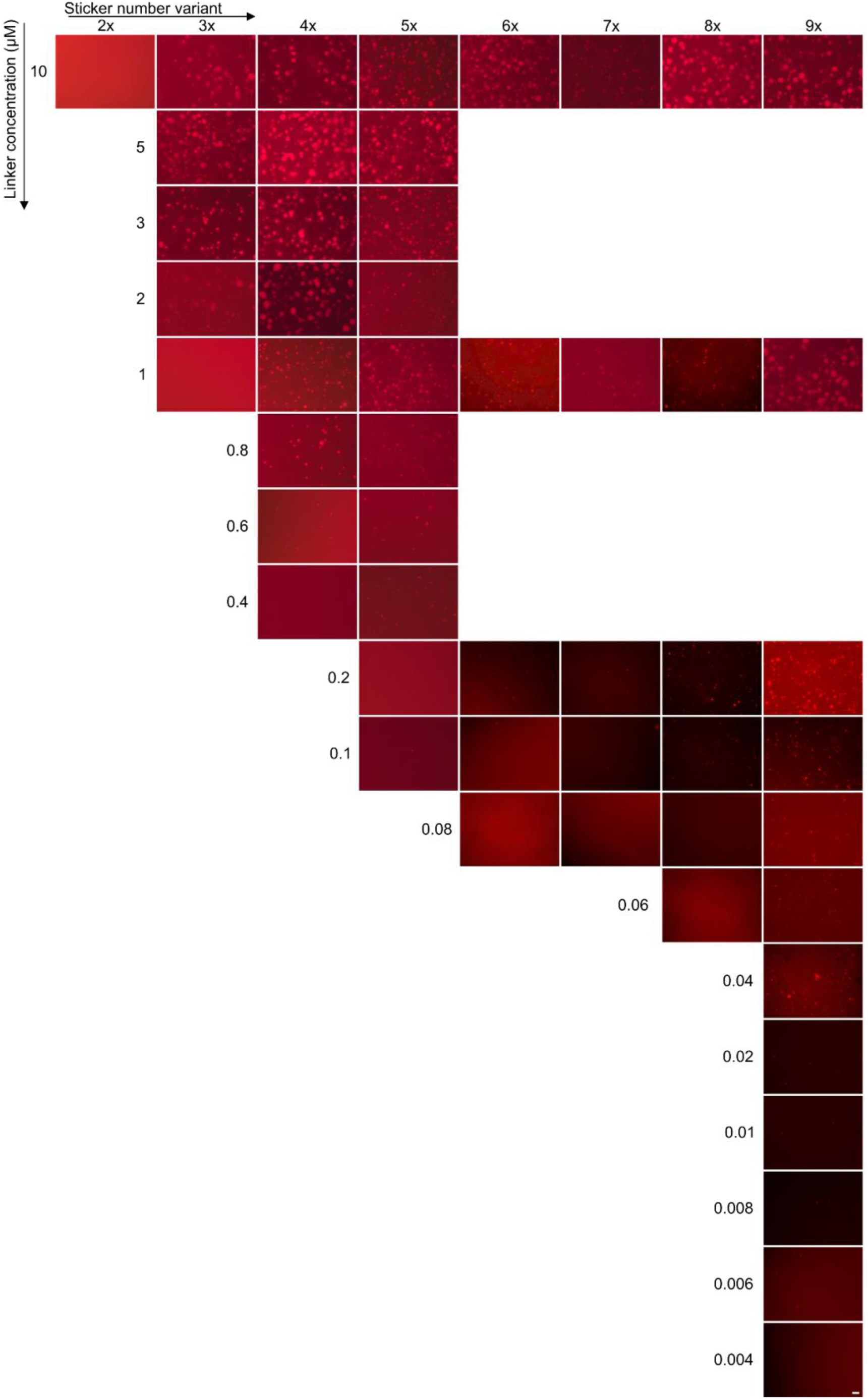
*In vitro* phase separation of EPYC1 sticker variants. 1 µM CrRubisco (0.5% Atto-594 labelled) mixed with the indicated concentrations of EPYC1 sticker variants. Scale bar = 10 µm.

**Fig. S4:**
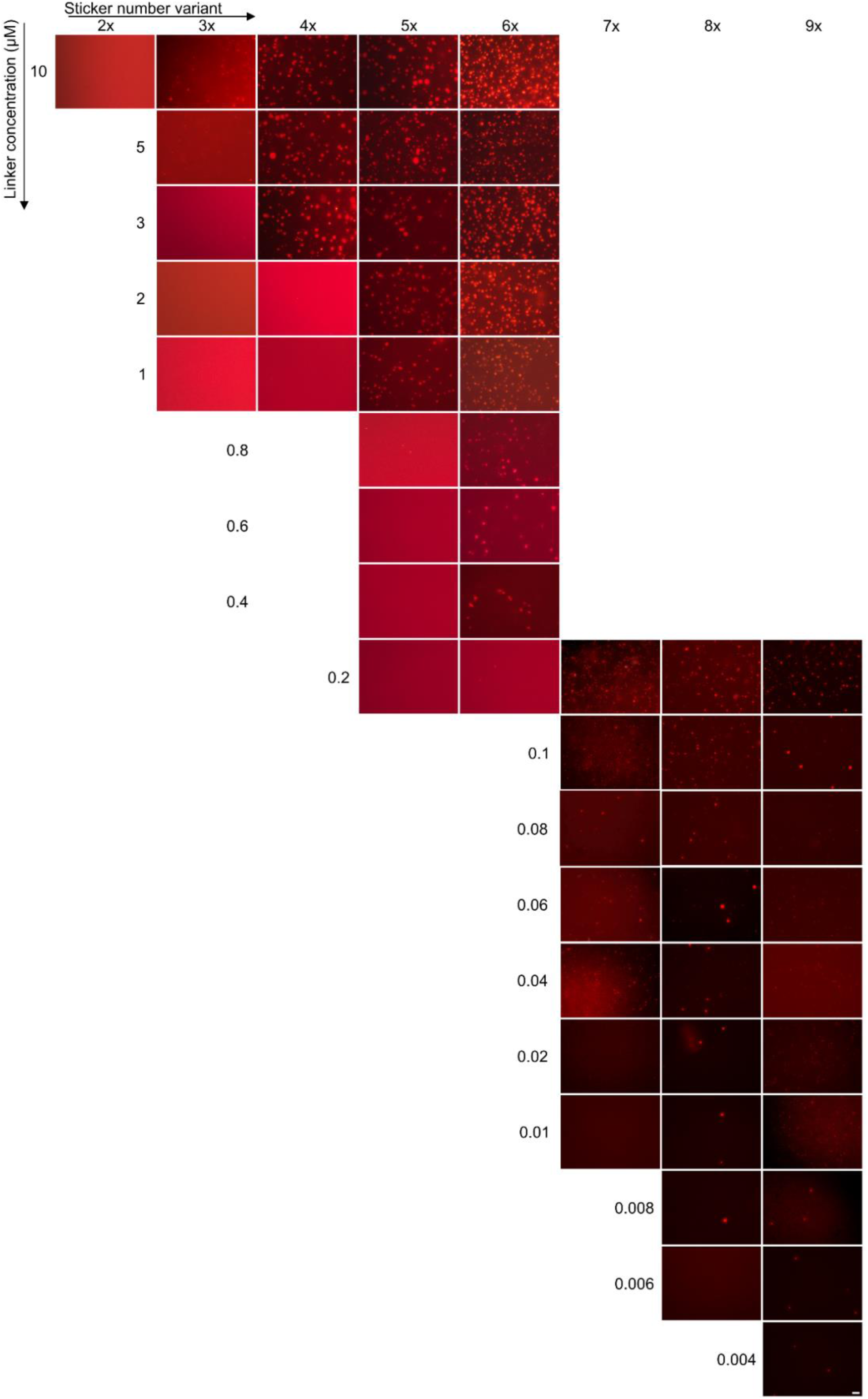
*In vitro* phase separation of CsLinker sticker variants. 1 µM CrRubisco (0.5% Atto-594 labelled) mixed with the indicated concentrations of CsLinker sticker variants. Scale bar = 10 µm.

**Fig. S5:**
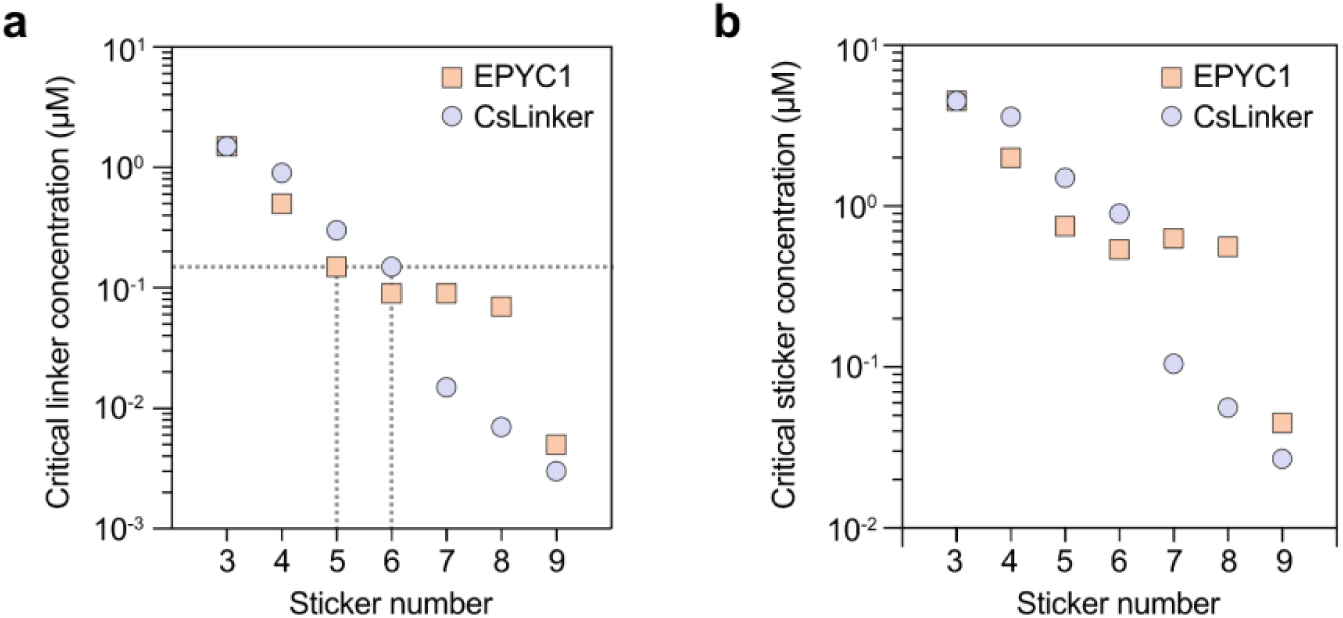
Sticker number reduces the critical concentration for *in vitro* Rubisco phase separation. **a)** Critical concentrations plotted as a function of total Linker protein concentration, derived from data in Fig. S2 and S3. Dotted lines indicate the critical concentration corresponding to the WT sticker numbers (5 for EPYC1; 6 for CsLinker). **b)** Critical concentrations normalised by total sticker concentration.

**Fig. S6:**
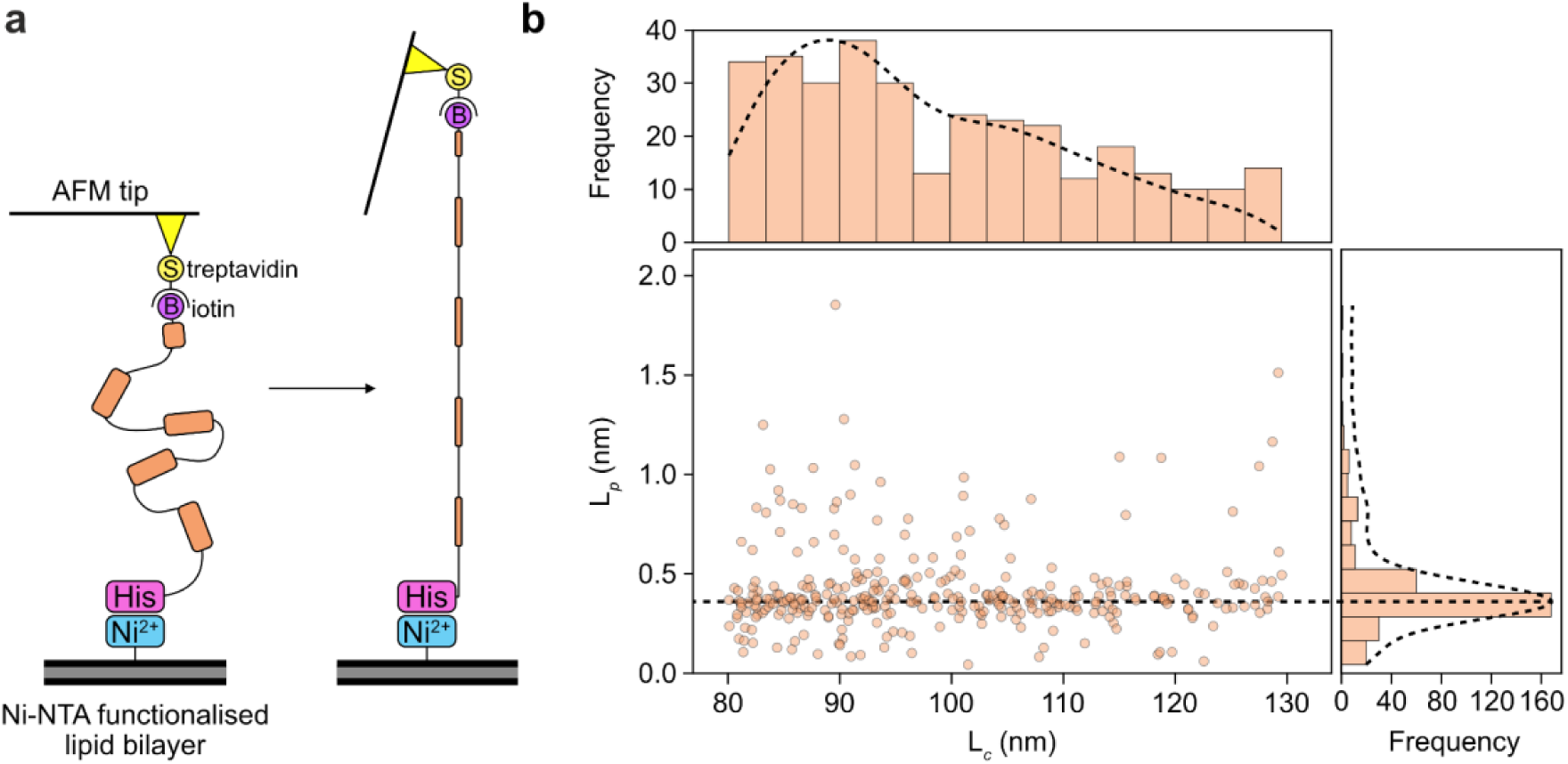
Atomic force spectroscopy measurements of EPYC1 persistence length (L*_p_*). **a)** Schematic of the tethered single-molecule AFM setup. EPYC1 is biotinylated at its C-terminus and coupled to the AFM cantilever via streptavidin, while the N-terminal 6×His tag is anchored to a Ni–NTA–functionalised supported lipid bilayer. **b)** Distributions of measured contour lengths (L*_c_*) and persistence lengths (L*_p_*). Only traces with contour lengths within 25% of the expected full length of EPYC1 were used to determine the consensus L*_p_*. Dotted lines indicate Gaussian fits to the binned data, and the resulting consensus L*_p_* is indicated.

**Fig. S7:**
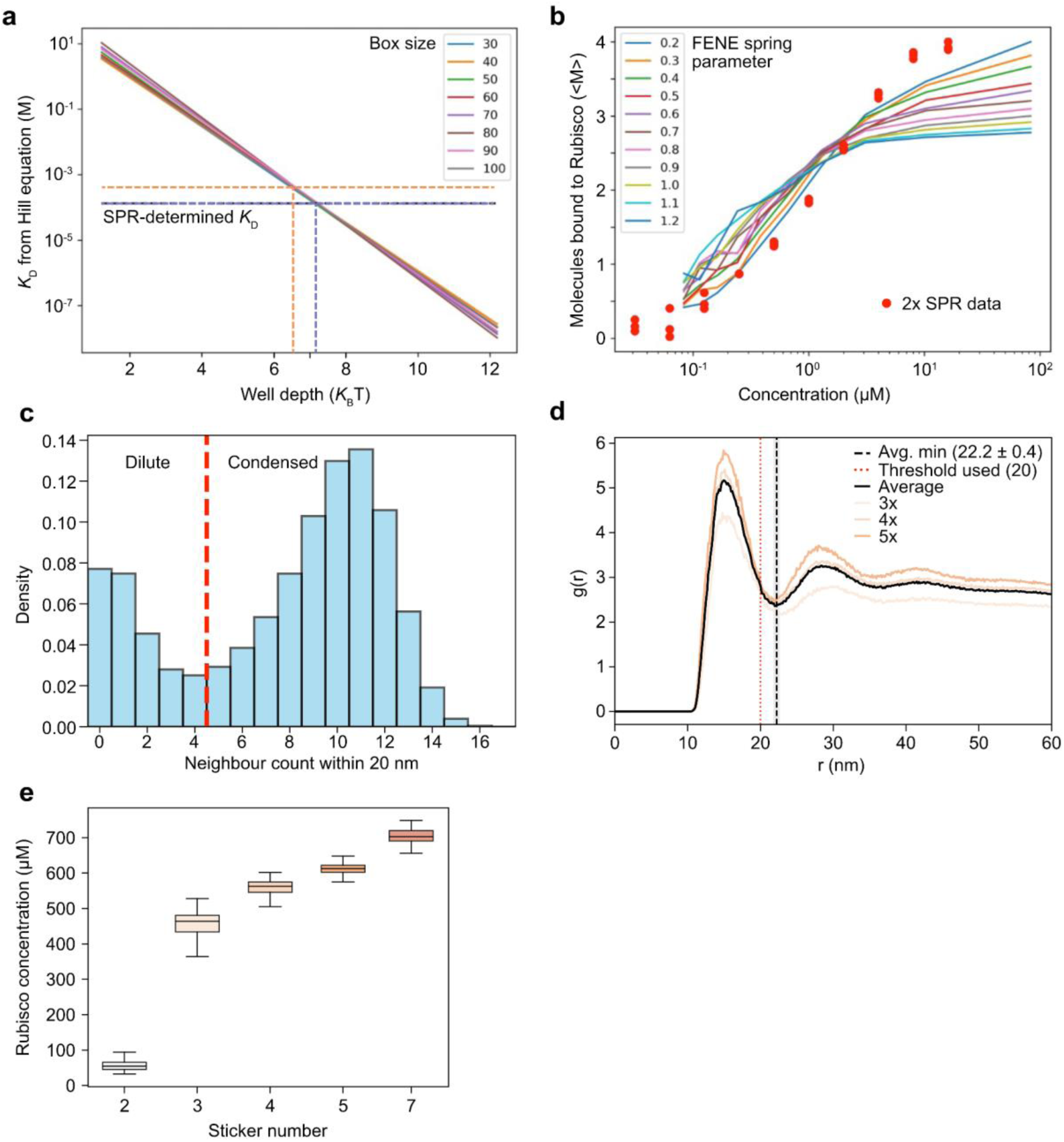
LAMMPS parameterisation and analysis. **a)** Relationship between the Lennard–Jones well depth used to model Linker–Rubisco interactions and the corresponding fitted dissociation constant *K*_D_. Dashed lines indicate experimentally measured *K*_D_ values for single sticker fragments of EPYC1 (orange) and CsLinker (purple) binding to their cognate Rubiscos, as reported by^13,27^. **b)** Determination of the FENE spring constant by fitting simulated force–extension behaviour to surface plasmon resonance (SPR) measurements of the 2x CsLinker construct. **c)** Histogram of Rubisco neighbour counts within a 20 nm radius, used to define the threshold for identifying condensed Rubisco in simulations. The threshold used to determine the number of neighbours to define Rubisco as being in the condensed phase is shown. **d)** Radial distribution function (RDF) plot of condensed Rubiscos in the 1:5 ratio simulations for 3x, 4x and 5x variants of EPYC1. The fitted minima between peak 1 is shown, as well as the threshold used in the calculations of partitioning (20 nm). **e)** Rubisco concentration in the condensed phase of LAMPPS simulations. Boxplot indicate median, IQR and range over last 200 frames of simulations.

**Fig. S8:**
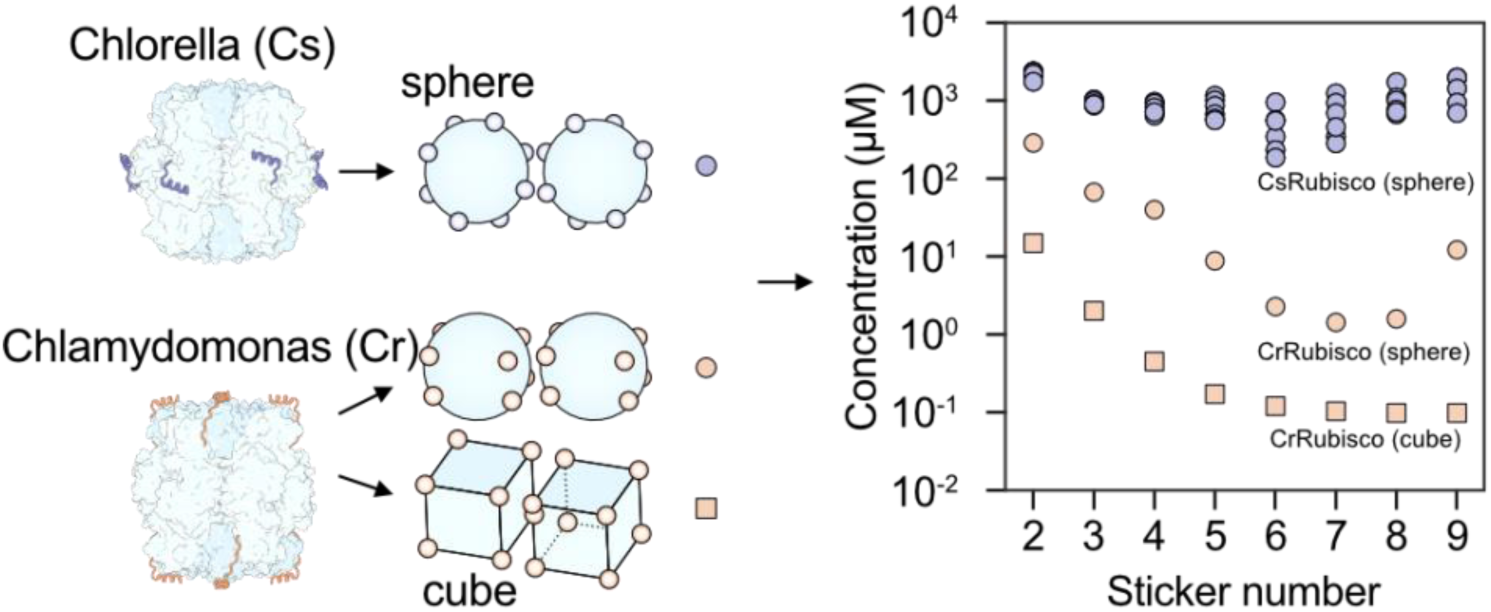
Effect of sticker binding positions on the critical total protein concentration required for phase separation across different Rubisco-Linker models. CrRubisco (cube), CrRubisco (sphere), and CsRubisco (sphere). For CsRubisco, no unique fit to the SPR data was obtained, so symbols indicate the range of values from the fit error.

## Supplementary note

By applying a cubic CrRubisco geometry with binding sites located at the corners, we previously determined the binding energy to be 11.8 *k*_B_*T* and the spacer flexibility parameterised using a freely jointed chain model with a Kuhn length of 0.88 ± 0.02 nm^13^. While this cubic geometry generated dimerization concentrations as an encouragingly good proxy for the critical concentration for 3-5 stickers, we here use a spherical geometry, as this better facilitates the positions of the binding sites according to their experimentally determined three-dimensional coordinates for both *Chlamydomonas*^23^ and *Chlorella*^27^. A new analysis of our *Chlamydomonas* data using the sphere model now yielded energy of 12.5 *k*_B_*T* and a Kuhn length of 0.58 ± 0.01 nm (a smaller Kuhn length corresponds to a spacer that is more difficult to extend). Through new measurements of the Kuhn length using atomic force microscopy (AFM), we find values of 0.72 ± 0.09 nm S.D. (Fig. S6) that are comfortably within this 0.58-0.88 nm range. For *Chlorella*, there was no unique fit; however, we found binding energies ranging from −15.2 to −11.9 *k*_B_*T* for a Kuhn length in the physically relevant range of 0.5 to 1 nm.

The dimerization concentration using the spherical Rubisco shape remains a reasonable proxy for the phase behaviour of CrRubisco, but turns out to be poor for CsRubisco (Fig. S8). Indeed, the dimerization concentration of CrRubisco decreases with an increasing number of stickers as before, albeit that the concentrations are shifted ∼30-fold due to increased excluded-volume interactions; this corresponds to an effective repulsion of ∼*k*_B_*T* ln(30) ∼ 3.4 *k*_B_*T* that could easily be compensated by weak non-specific interaction between the Rubisco with each other and/or the Linker protein. However, for *Chlorella* we found the dimerization concentration increased by another ∼10-1000-fold, which reflects the previous notion by GrandPre et al., 2023, that equatorial geometries are unfavourable for self-assembly. However, arguably the pairwise interaction is a poorer phase-separation predictor for CsRubisco than for CrRubisco, as the equatorial arrangement of the binding sites may require a larger number of Rubiscos to interact to form a stable cluster.

**Fig. S9:**
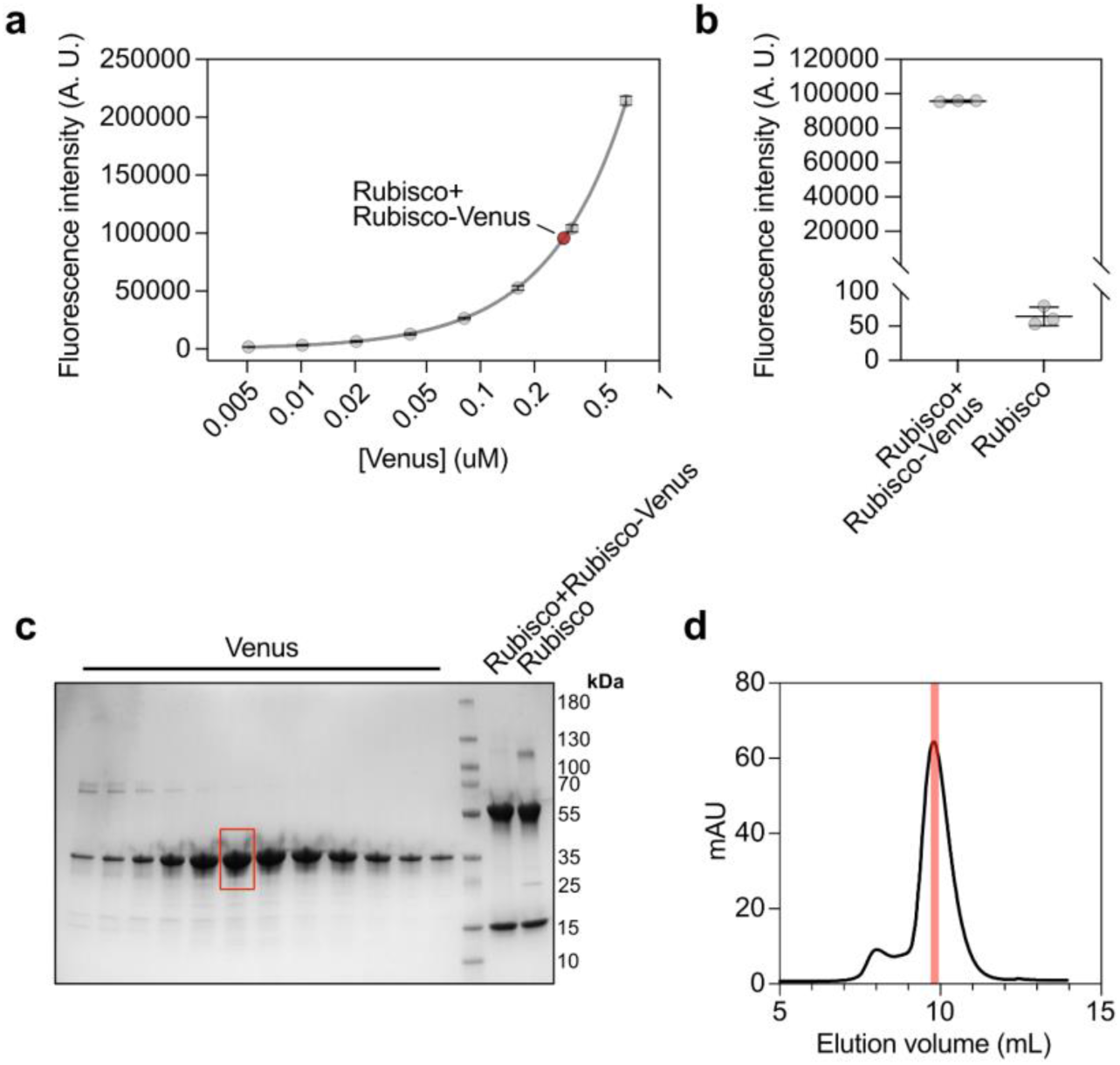
Quantification of RbcS-mVenus labelling proportion. **a)** Fluorescence intensity of the Rubisco / Rubisco-mVenus pool that was purified from the chassis line, relative to a standard curve of purified Venus. Rubisco was 3.85 µM in this measurement. **b)** Fluorescence intensity of the Rubisco / Rubisco-mVenus pool relative to the same concentration of untagged Rubisco purified from WT. **c)** SDS-PAGE analysis of purified proteins used for quantification. The fraction of purified Venus used for the quantification is highlighted by the red box. **d)** Size exclusion chromatography elution from Superdex 75 pg 10/300 GL column. The fraction used for quantification is shown in red.

**Fig. S10:**
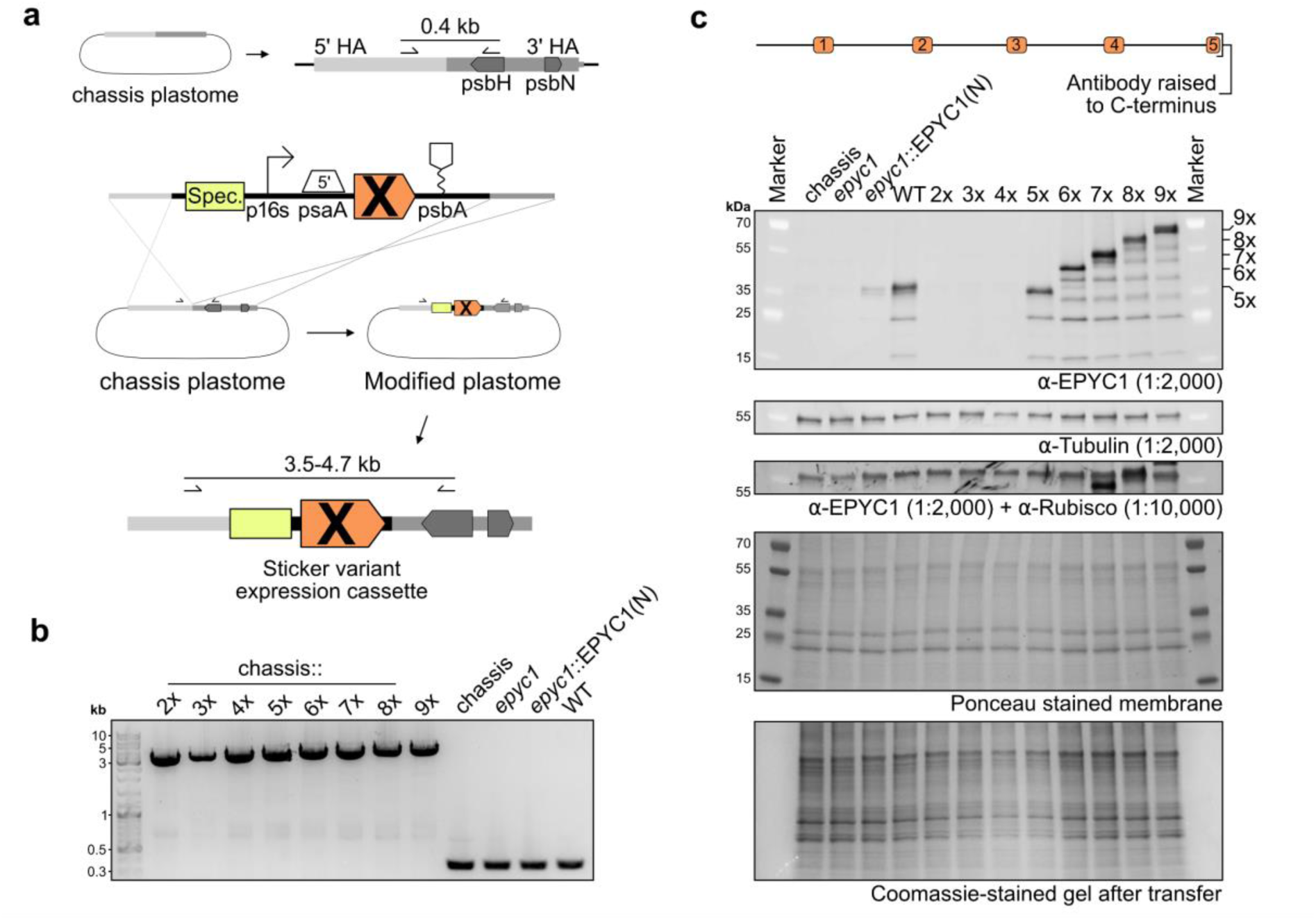
Validation of chloroplast expression lines. **a)** Schematic showing the positions of genotyping primers on the homology arms (HAs) of the chloroplast insertion cassette used to assess homoplasmy. The structure of the expression construct and its integration into the chassis plastome are shown. **b)** Genotyping PCR confirms homoplasmic integration of the EPYC1 sticker-variant constructs into the chloroplast genome (plastome) of the chassis line. epyc1::EPYC1(N) refers to the nuclear-expressed EPYC1 complementation strain from Mackinder et al. (2016). **c)** Western blot quantification of expression levels across strains. Because the EPYC1 antibody was raised against the C-terminus of EPYC1, variants lacking this region (shorter than WT; see Fig. S2) are not detected. Tubulin was used as a loading control.

**Fig. S11:**
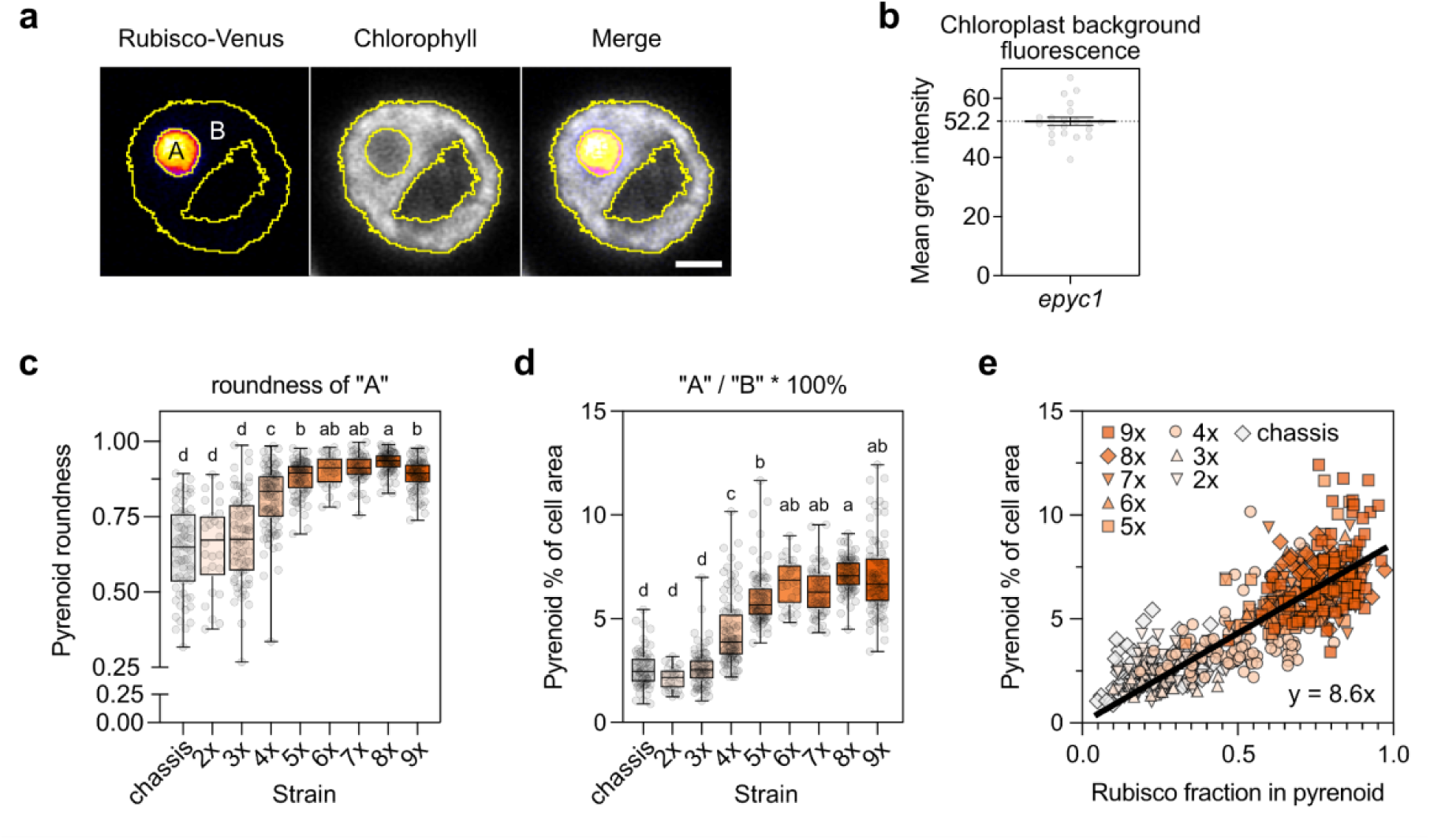
Quantification of pyrenoid size and shape across sticker-number variants. **a)** Example ROIs illustrating the segmentation workflow for Rubisco partitioning, showing pyrenoid, chloroplast, and whole-cell masks. Scale bar = 2 µm. **b) B**ackground fluorescence intensity measured in the epyc1 knockout strain, used for background subtraction during partitioning analysis. Error bars indicate s.e.m. **c)** Roundness of segmented pyrenoid cross-sections for each strain. Box plots show the median (line), mean (cross), interquartile ranges (box), and whisker limits (1.5x IQR); statistical groups reflect *p* < 0.001. **b)** Percentage of total cell cross-sectional area occupied by the pyrenoid. Box plots and statistical conventions as in panel a. **c)** Relationship between the fraction of cellular Rubisco localized to the pyrenoid and the pyrenoid cross-sectional area, with a linear fit across all strains.

**Fig. S12.**
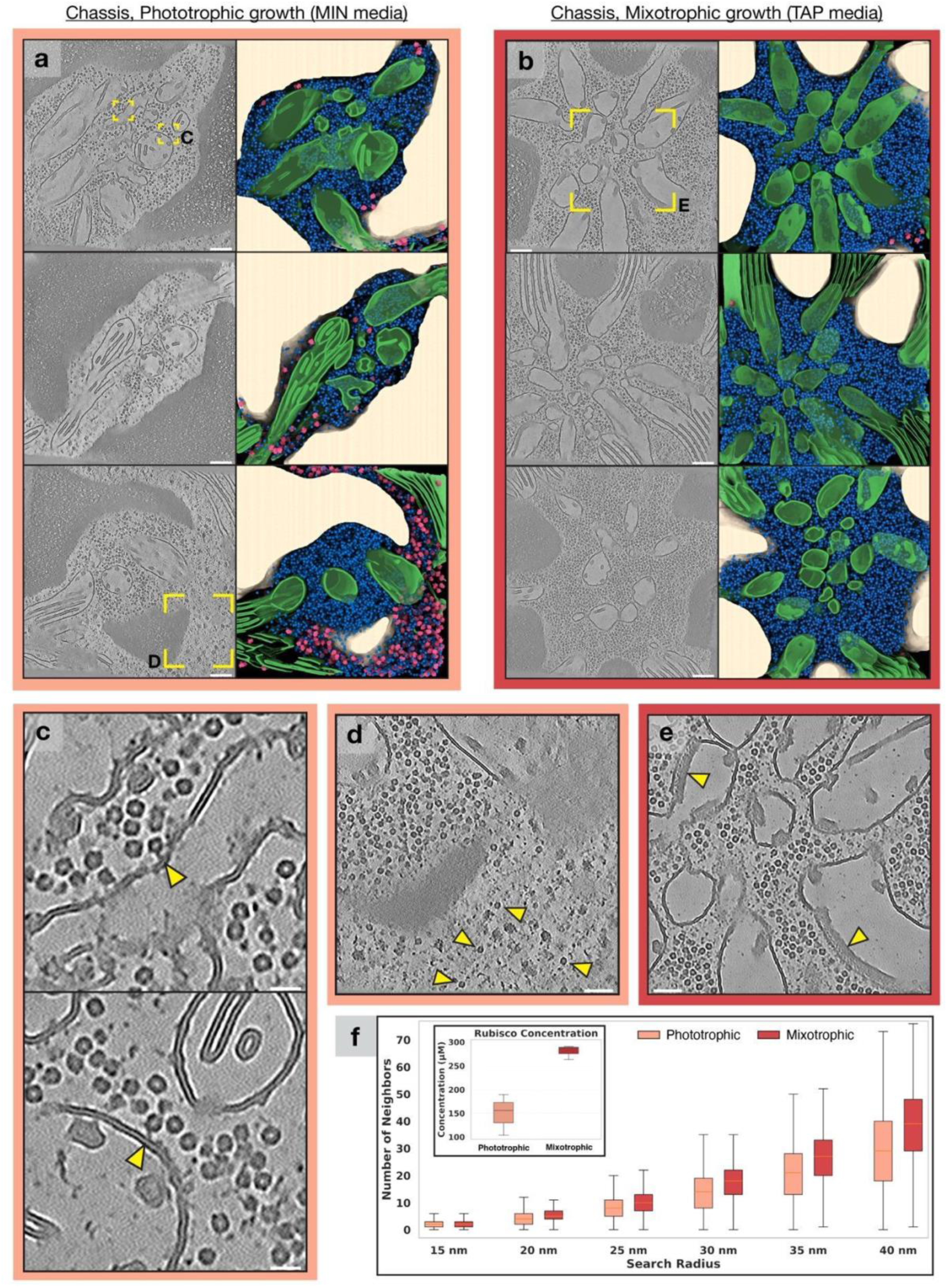
Cryo-ET of the chassis strain grown under autotrophic vs. mixotrophic conditions. **a)** Three representative tomograms and segmentations of chassis pyrenoids in phototrophic conditions. Rubisco is shown in blue, thylakoids and pyrenoid tubules in green, and chloroplastic ribosomes in pink. **b)** As in a) but grown in mixotrophic conditions. Scale bars, 100 nm. **c)** Zoom in showing two cases of Rubisco tethered to the tubule membrane (yellow arrows). Scale bar, 20 nm. **d)** Zoom in showing the partial loss of a condensed pyrenoid matrix. Yellow arrows indicating Rubisco’s intermixed with ribosomes in the stroma. Scale bar, 50 nm. **e)** Yellow arrows showing an unknown protein array decorating the minitubule membrane upon fusion with the tubule membrane. Scale bar, 50 nm. **f)** Box plot showing the number of neighbours for each Rubisco particle within a given spherical search radius in the two conditions. Inset showing the difference in Rubisco concentration (particles per volume extrapolated to micromolar) between the two conditions.

**Fig. S13.**
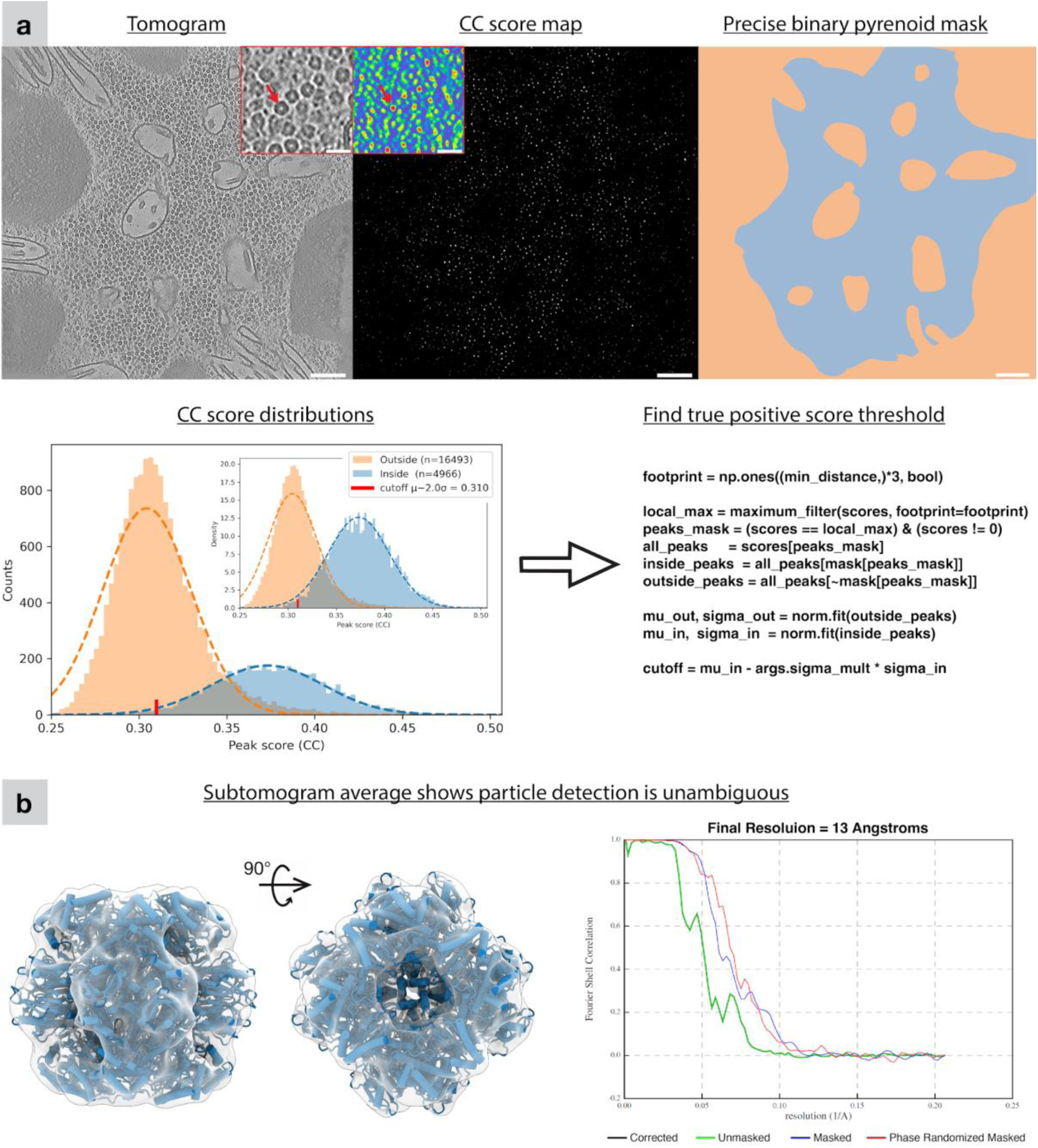
Cryo-ET processing pipeline. **a)** Top row, from left to right: tomogram, cross-correlation (CC) score map from template matching, and precise binary pyrenoid mask used for volume estimation and masking of the score map. Scale bar, 100 nm. Insets show close up Rubisco’s and their high CC scores as a heat map (red arrows). Scale bars, 20 nm. Bottom left panel: CC distributions with Gaussian fits of the masked area where inside is the true positive containing pyrenoid matrix and outside is false positive containing non-pyrenoid area (e.g. starch and tubules). The CC cutoff was defined as the median of the inside distribution minus two standard deviations of its fitted Gaussian. The inset shows the same data as a normalized probability distribution rather than raw counts. The arrow highlights Python-style pseudocode implementing the peak-detection and CC thresholding procedure. **b)** Map from subtomogram averaging of all positions and orientations determined by template matching and used for the spatial organisation analyses. A cartoon-style model of Rubisco (PDB 7JN4) is fitted in blue using ChimeraX. The gold-standard Fourier shell correlation (FSC) between two half-maps (RELION5) indicates a resolution of 13 Å at the 0.143 cutoff.

**Fig. S14:**
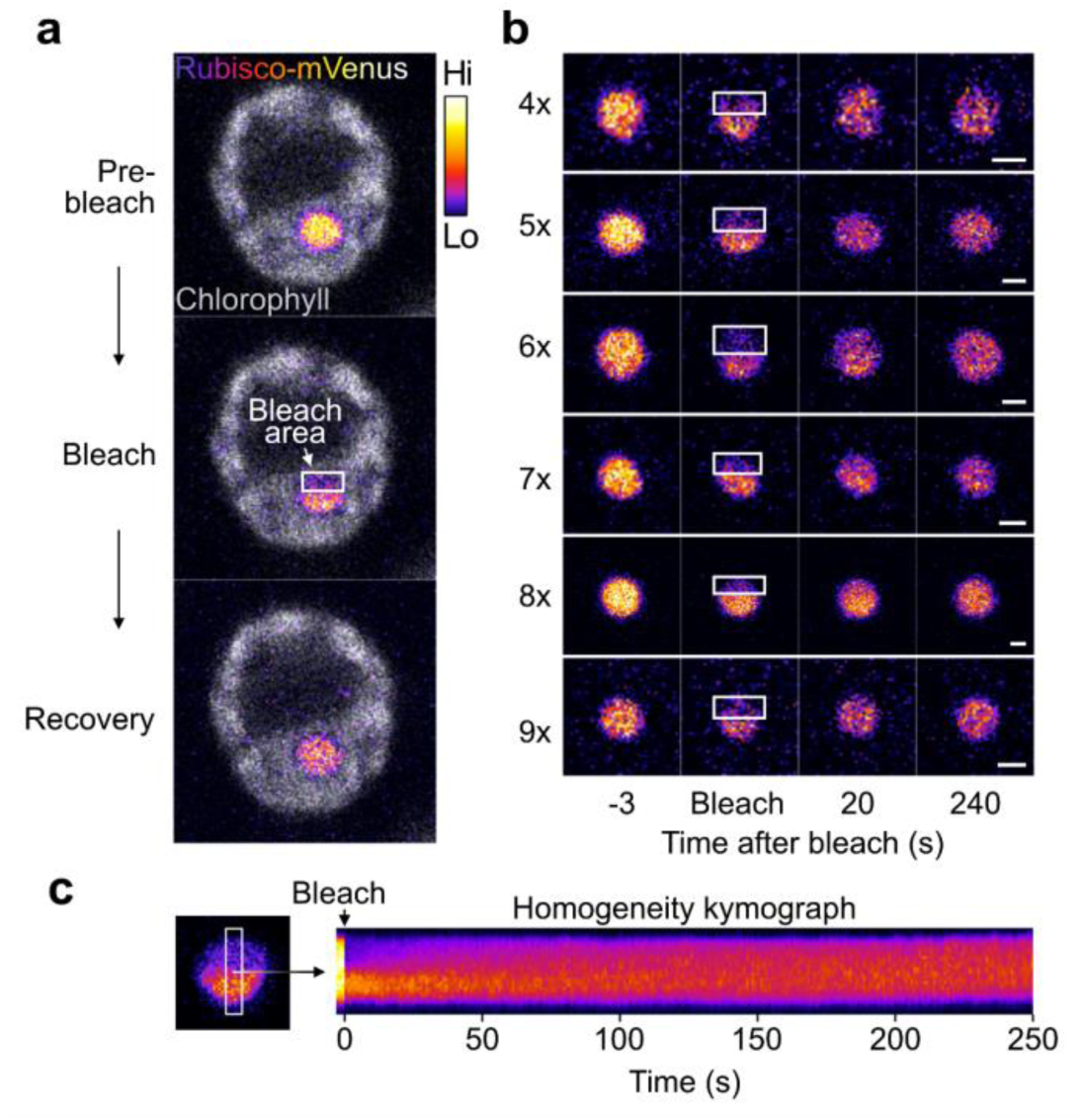
Fluorescence recovery after photobleaching (FRAP) of Rubisco in Chlamydomonas pyrenoids. **a)** Representative snapshots from a FRAP experiment showing a single pyrenoid. Chlorophyll autofluorescence is shown in grayscale, and Rubisco-mVenus signal is displayed using a fire color scale (see inset key). The bleached region is indicated. **b)** Examples snapshots of recovery during experiments with each of the tested variants. **c)** Example kymograph of Rubisco-mVenus signal across the bleached region over time, demonstrating recovery and redistribution of Rubisco within the pyrenoid matrix.

**Fig. S15:**
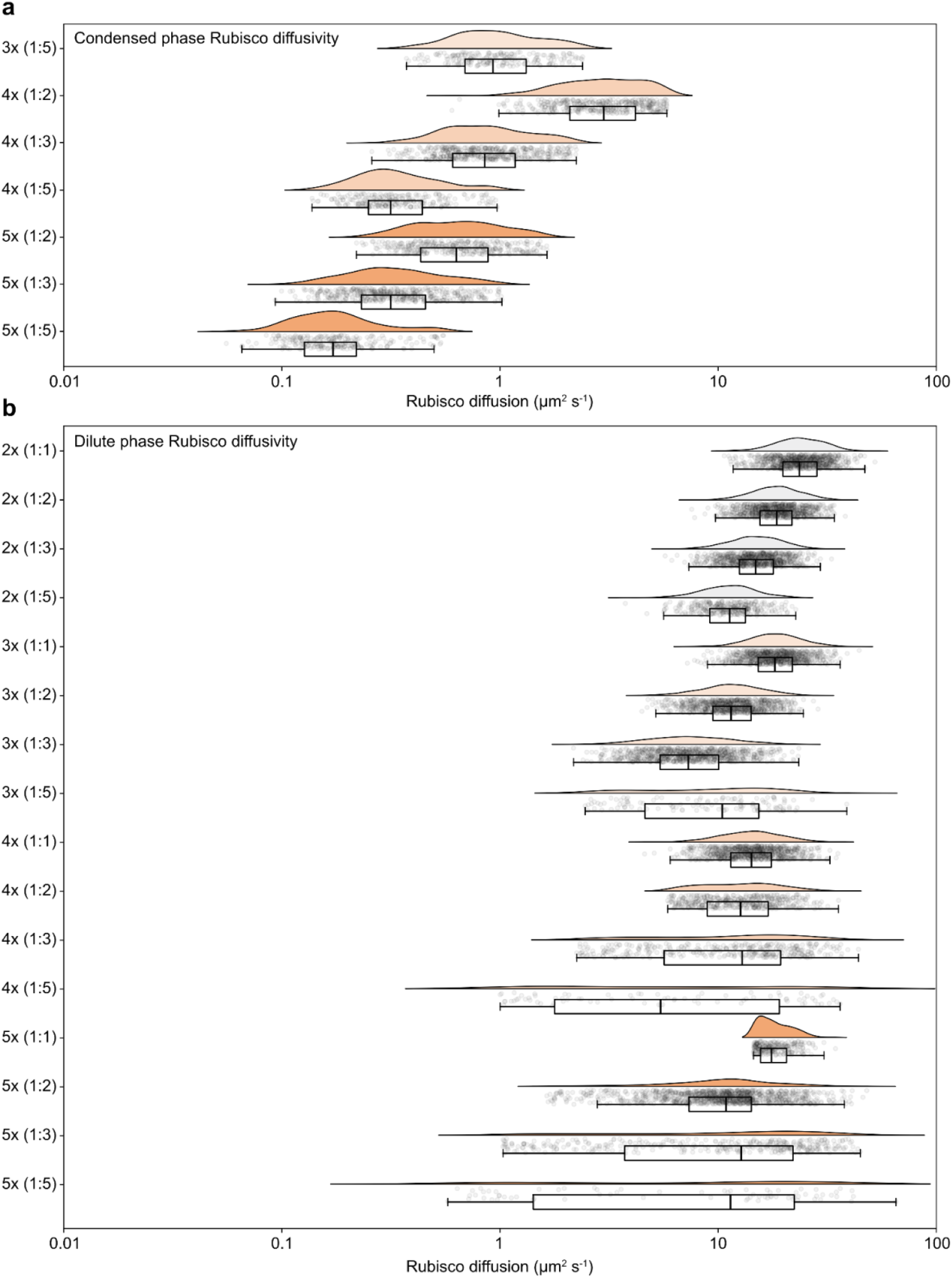
Diffusivity of Rubisco in LAMMPS simulations with EPYC1 variants. **a)** Mean-squared displacement–derived diffusivity of Rubisco molecules within the condensed phase at the indicated Rubisco:Linker concentrations and sticker variants. **b)** Corresponding diffusivity of Rubisco in the dilute phase. Box plots indicate the median (line), mean (cross), interquartile range (box), and whiskers extending to 1.5x the IQR.

**Fig. S16:**
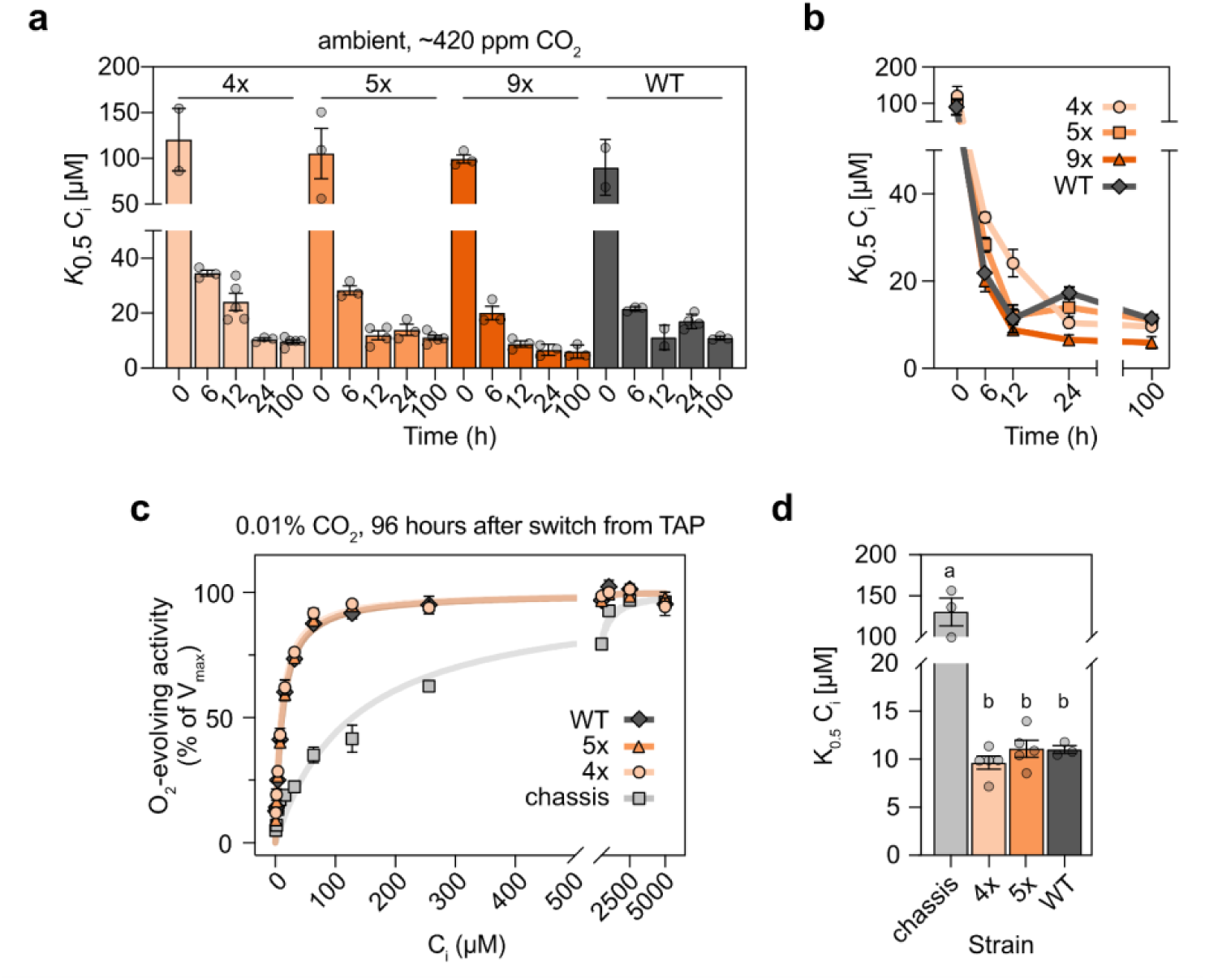
Photosynthetic efficiency of sticker variant lines relative to WT. **a)** Affinity for inorganic carbon (*K*_0.5_ C_i_) measured by oxygen evolution. Mean ± s.e.m. are indicated. **b)** Plot of mean *K*_0.5_ C_i_ against time, where error bars indicate s.e.m. **c)** Oxygen evolution rate as a percentage of the fitted V_max_ as a function of inorganic carbon (C_i_) concentration, measured 96 hours after the switch from mixotrophic to phototrophic growth conditions under which the CO_2_ concentration was 0.01%. For chassis, 4x, 5x and WT, *n* = 3, 5, 5 and 3 respectively. The Michaeles-Menten fit to the average of the replicates is shown. **d)** Fitted K_0.5_ C_i_ values from curves in (c). Statistical groups indicate *p* < 0.001.

**Fig. S17:**
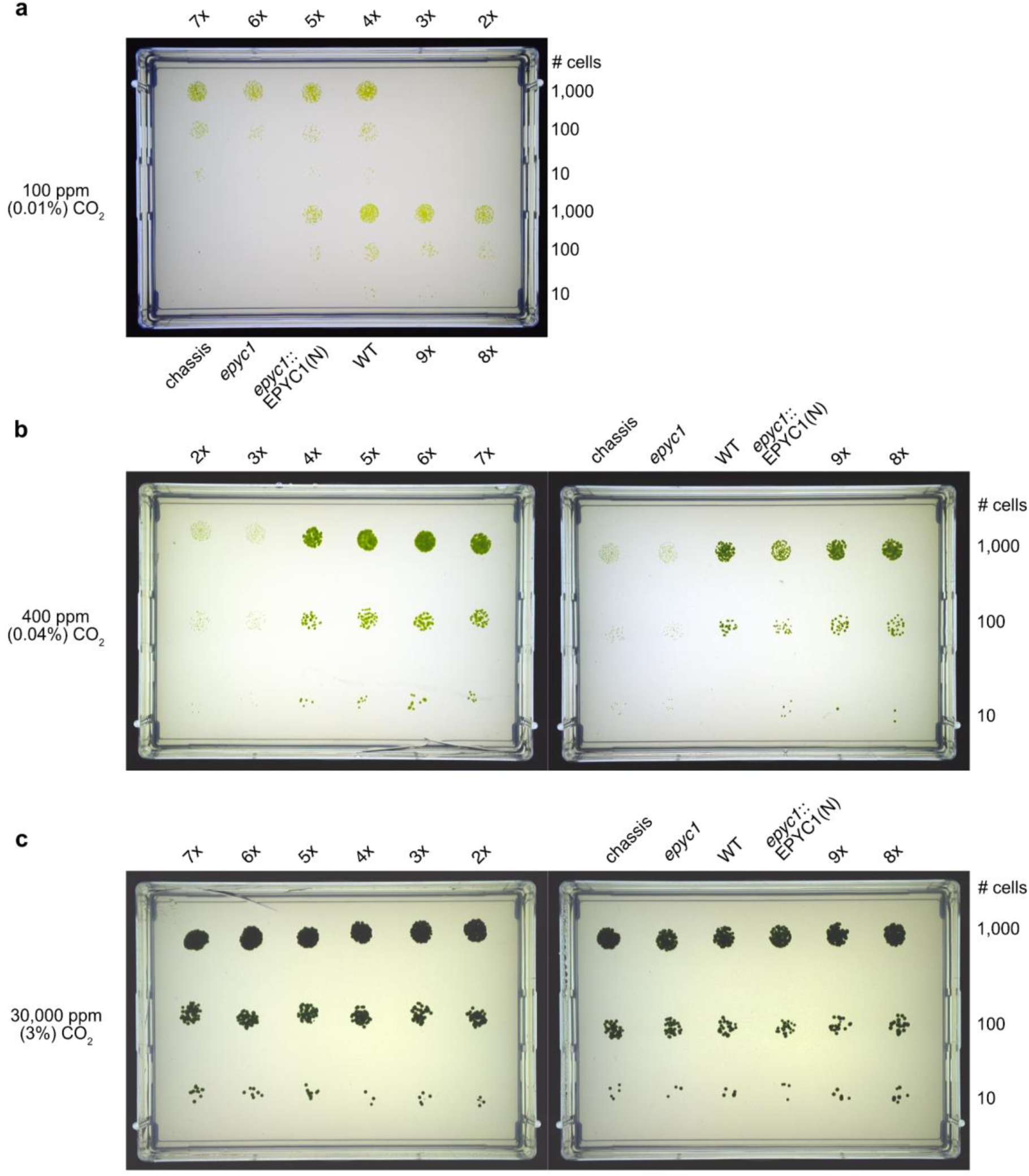
Growth phenotypes of EPYC1 sticker-variant lines under different CO_2_ conditions. **a)** Uncropped spot assay showing growth of strains at 100 ppm CO_2_ (0.01%). **b)** Growth under ambient CO_2_ (400 ppm; 0.04%). **c)** Growth under elevated CO_2_ (30,000 ppm; 3%).

**Fig. S18:**
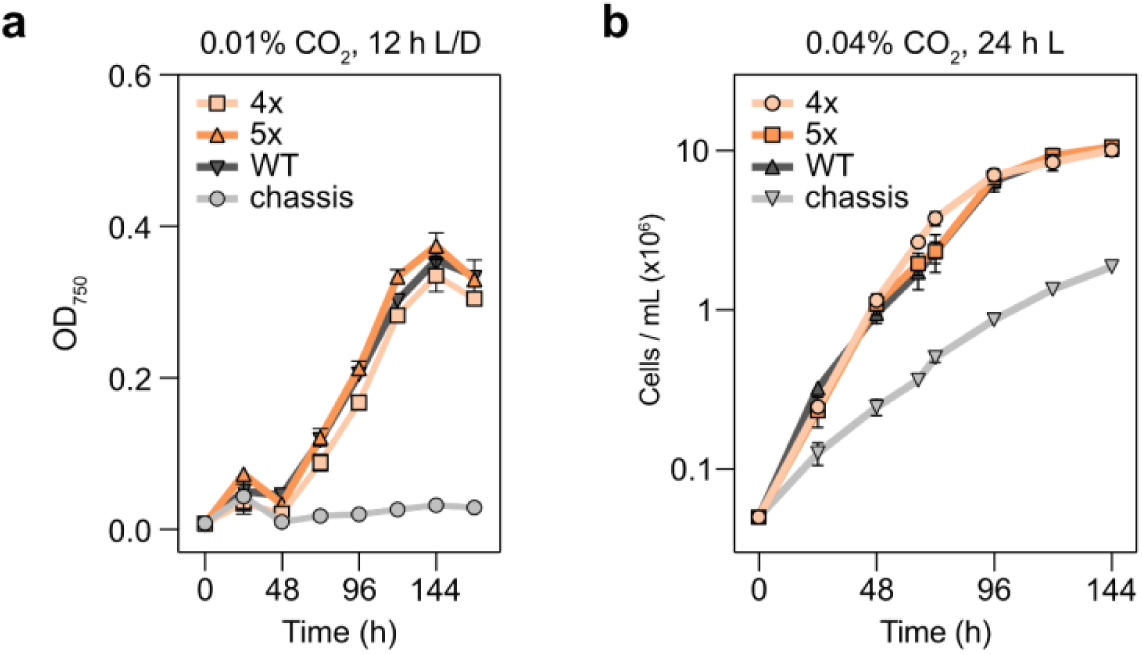
Liquid growth phenotypes of 4x and 5x lines. **a)** Optical density at 750 nm (OD_750_) of cultures grown under low CO_2_ (0.01%, 100 ppm) in 12 h light / 12 h dark diurnal cycles. **b)** Cell counts of cultures grown under ambient CO_2_ (∼0.04%, 420 ppm) in constant light (24 h). Error bars indicate s.e.m.

**Fig. S19.**
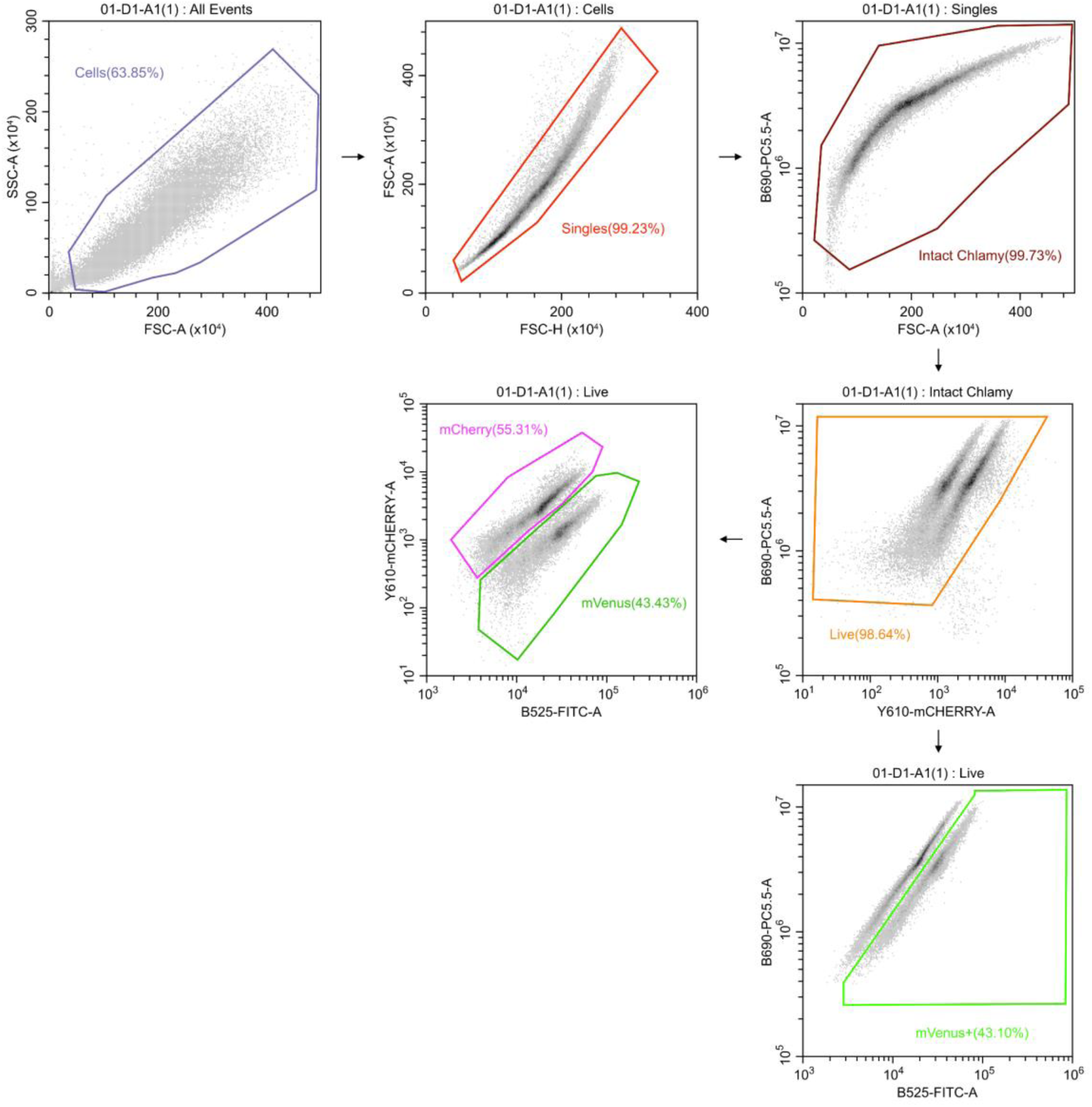
Gating workflow for flow cytometry measurements. For both dual-colour and single-colour experiments, initial gating steps were performed identically (up to the “intact Chlamy” gate). For single-colour experiments (*e.g.*, comparing 4x,mCherry to *epyc1* [untagged]), cells were subsequently gated based on mVenus and chlorophyll autofluorescence (bottom right). For dual-colour experiments, gating was performed on both fluorophore channels (mCherry and mVenus) to isolate the appropriate populations.

**Fig. S20:**
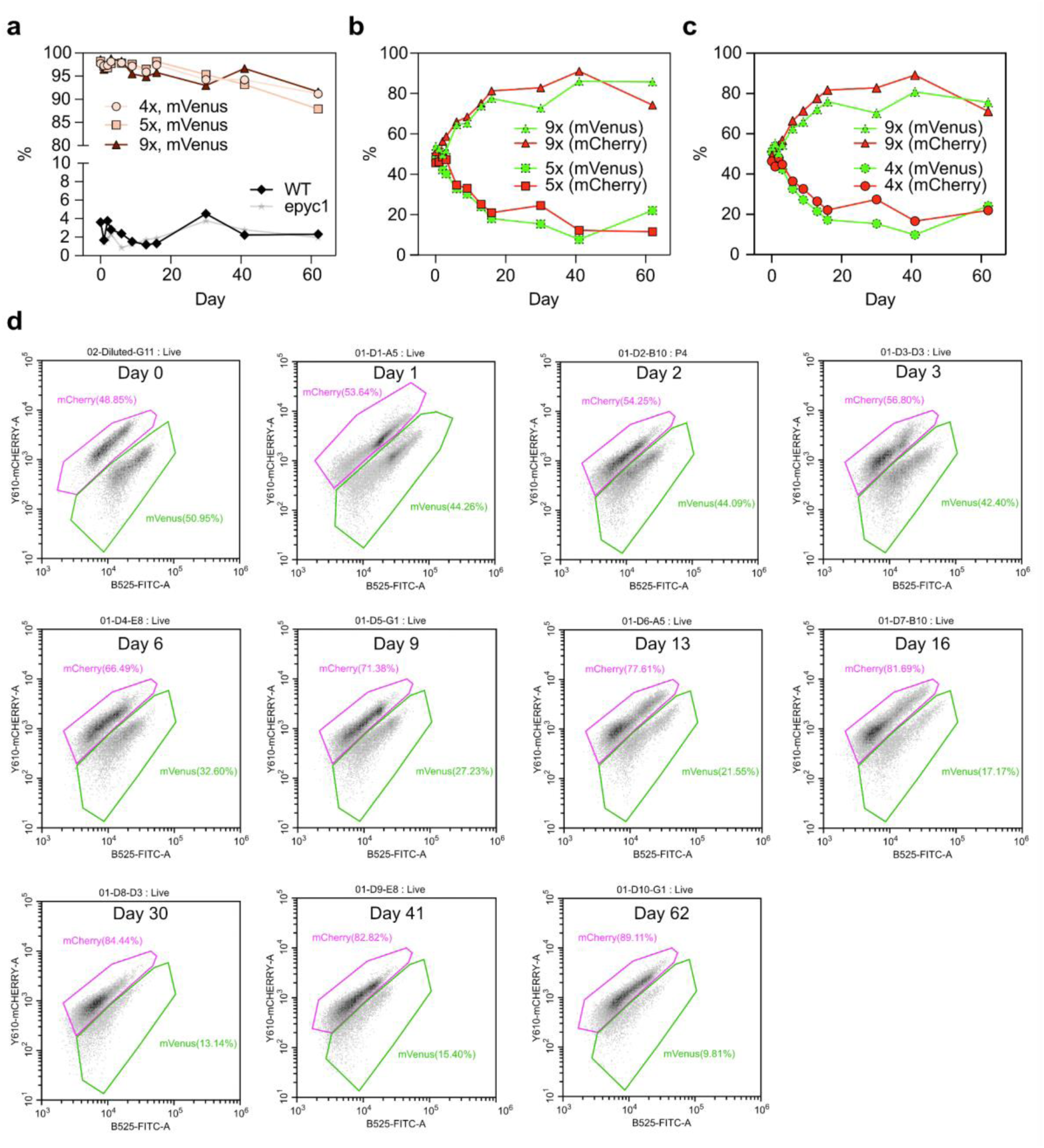
Additional flow cytometry analyses. **a)** Percentage of cells with mVenus fluorescence above the defined gating threshold across the duration of the experiment. Expression lines show a gradual decline in reporter signal, while untagged controls (WT, *epyc1*) show no increase in background fluorescence. **b)** Replicate traces from competitive growth assays comparing 5x and 9x variants, demonstrating strong agreement between reciprocal colour-swap experiments. **c)** Replicate traces from competitive growth assays comparing 4x and 9x variants. **d)** Representative flow cytometry density plots from the 4x (mVenus) versus 9x (mCherry) competitive growth assay over time, illustrating the progressive population shift toward the 9x, mCherry-expressing line.

**Fig. S21:**
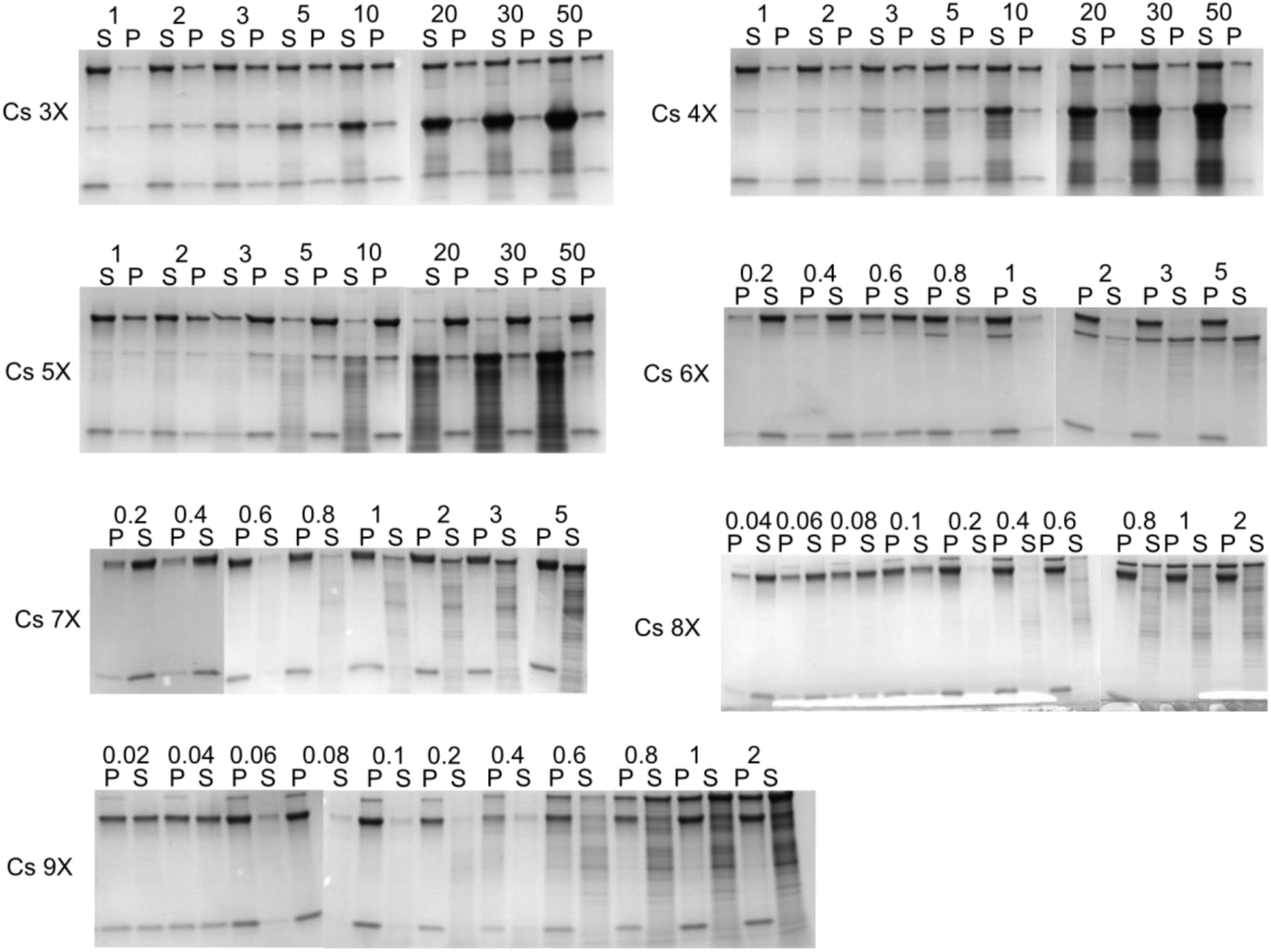
Droplet sedimentation assay gel images. SDS-PAGE gel images quantified in Fig. 1h.

**Table S1:**
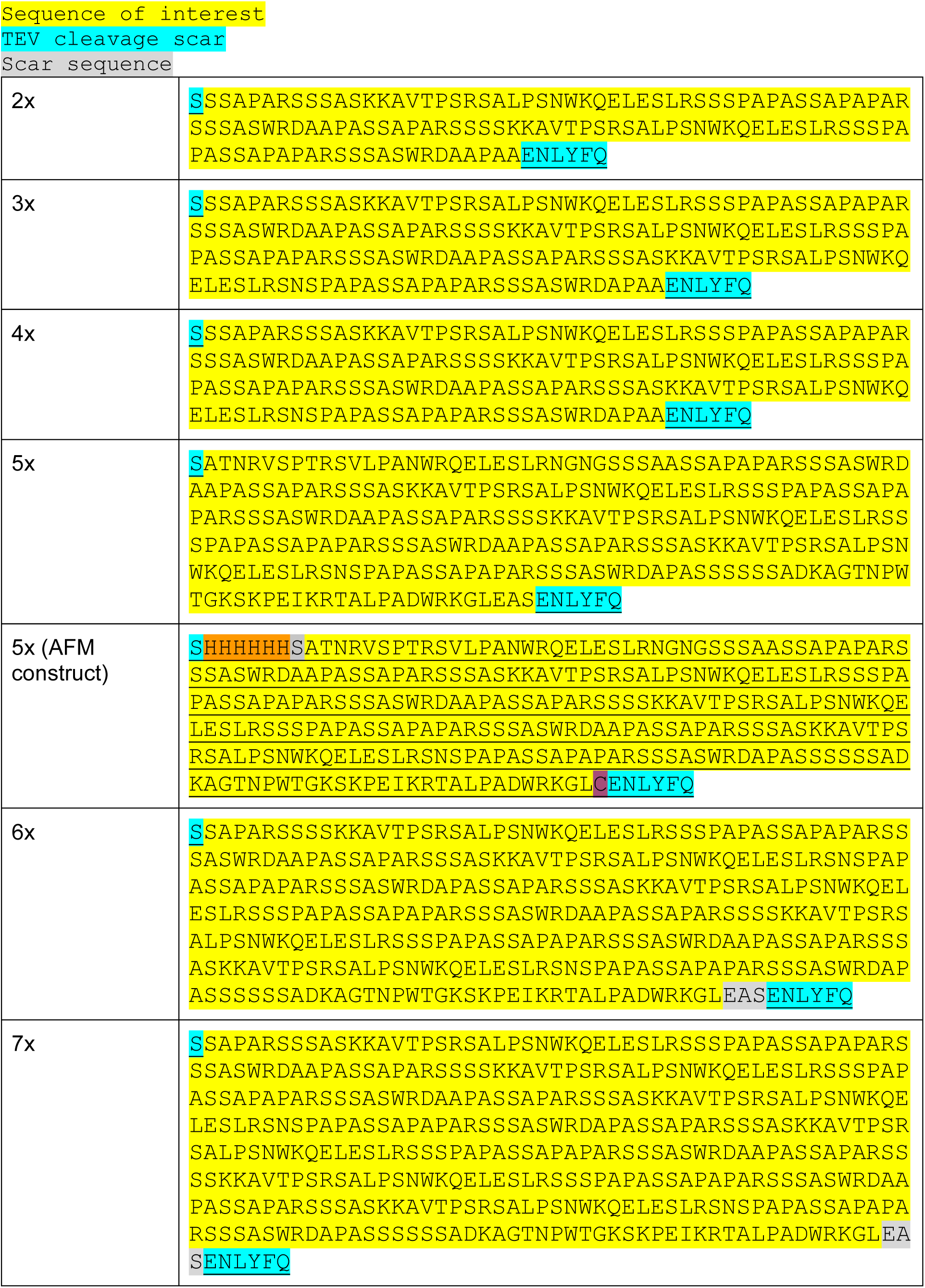
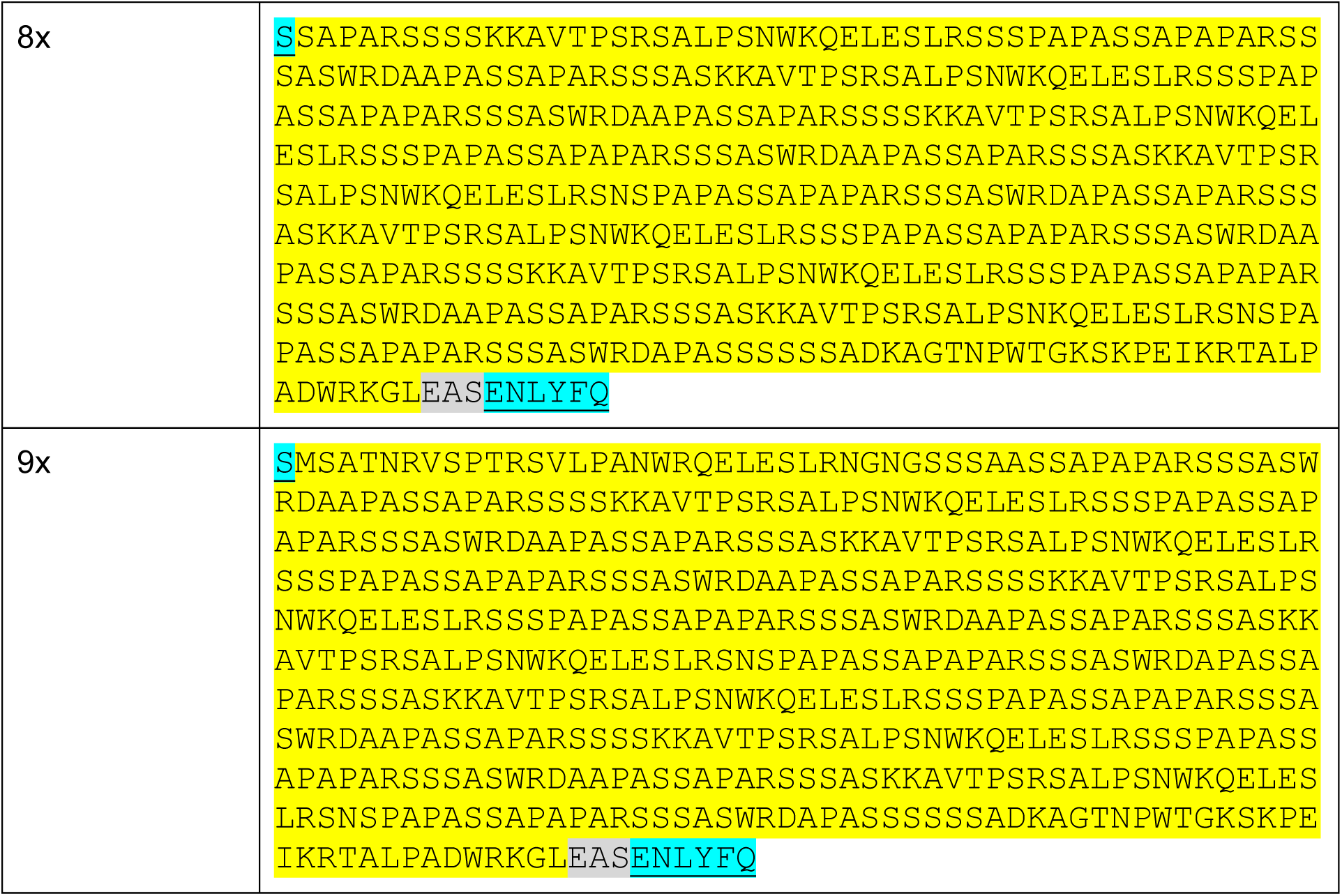
EPYC1 protein sequences used in *in vitro* experiments.

**Table S2:**
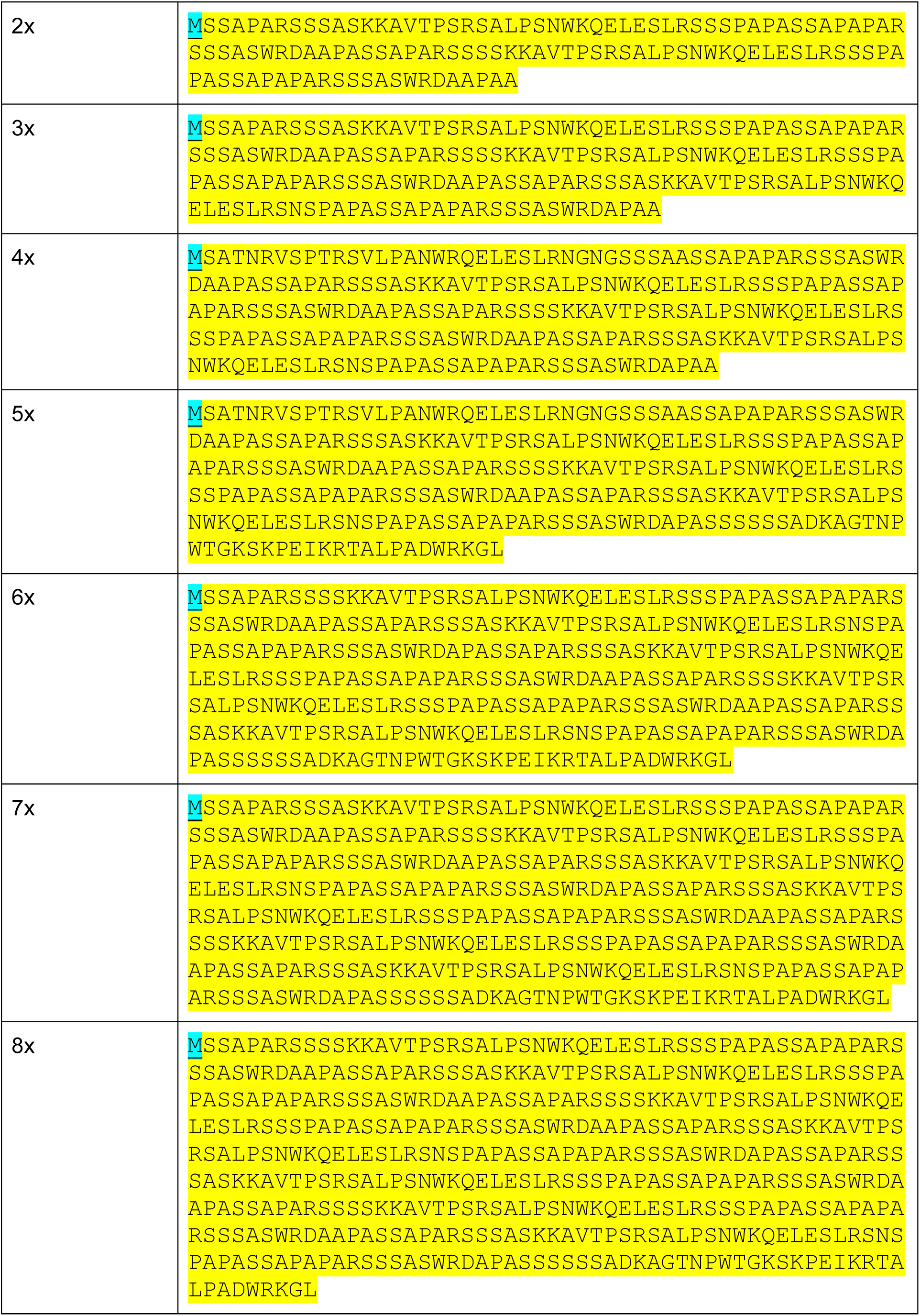
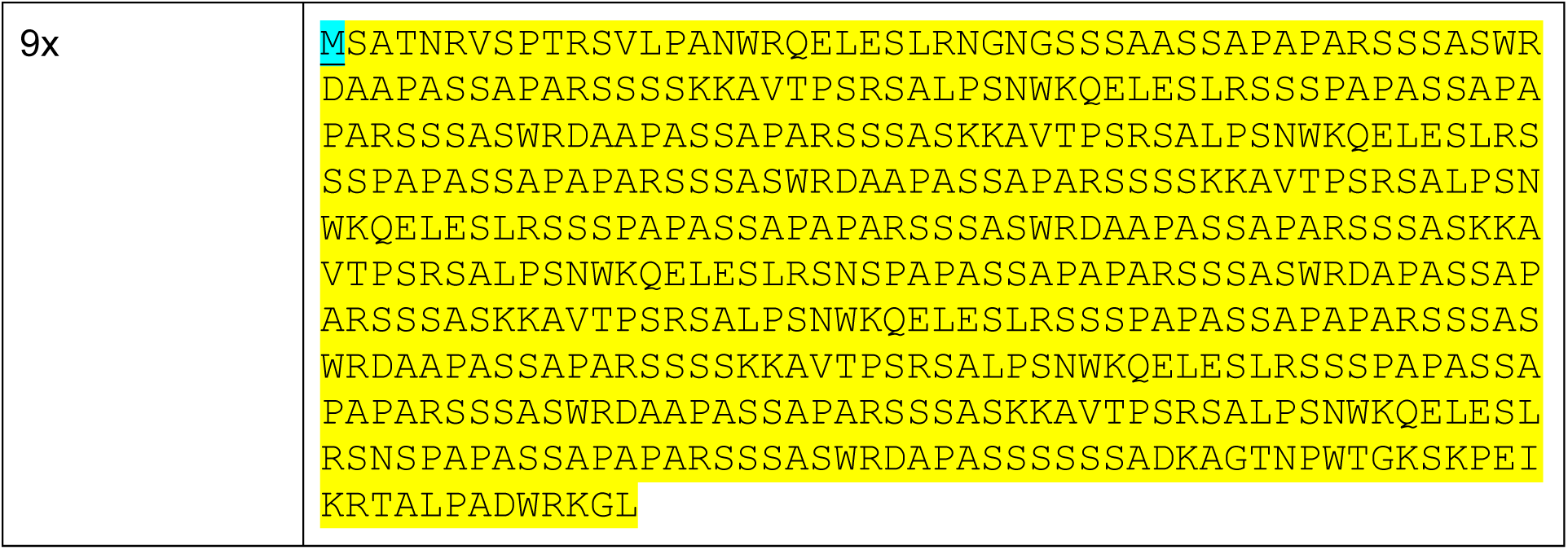
EPYC1 protein sequences used in *in vivo experiments*.

**Table S3:**
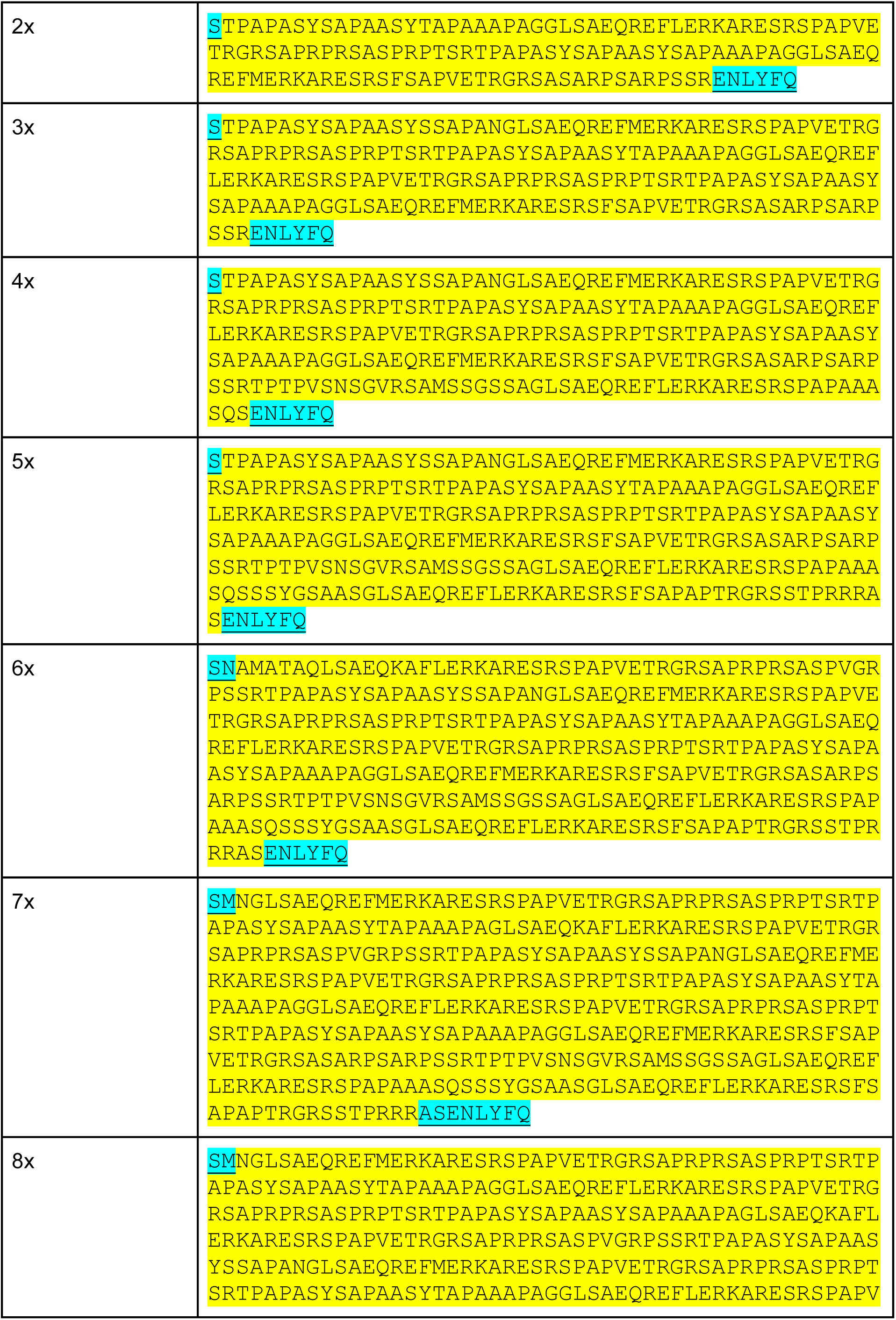
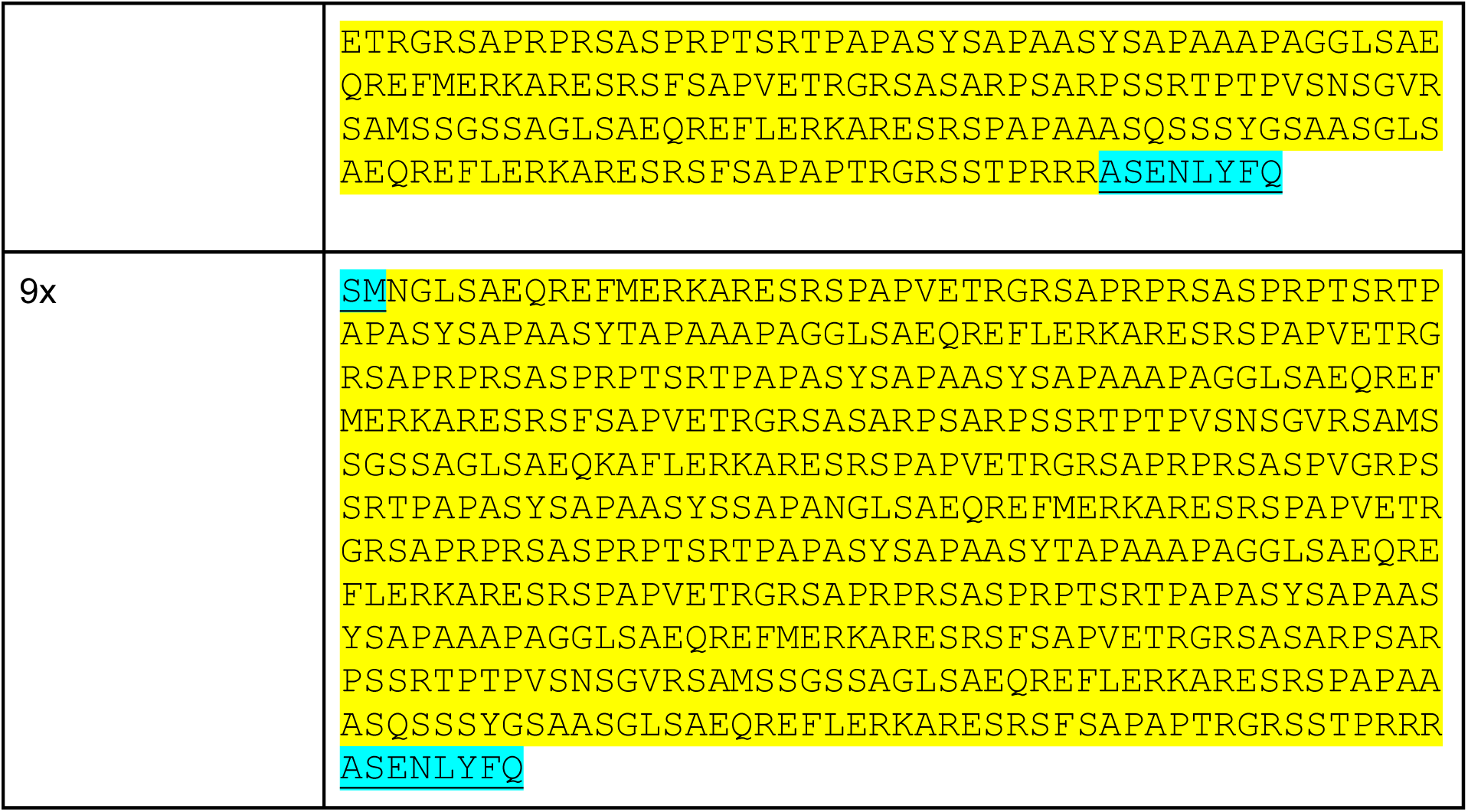
CsLinker protein sequences used in *in vitro* experiments.

